# The City Nature Challenge increases urban biodiversity knowledge and public engagement with blue spaces

**DOI:** 10.64898/2026.03.19.712856

**Authors:** Matthew C. Morgan, Charlotte R. Hopkins, Rodney Forster, Africa Gómez

## Abstract

Global biodiversity is declining at an unprecedented rate due to rapid environmental change and increasing human pressures. Ongoing urban expansion fragments natural systems, while urban design increasingly seeks to mitigate these impacts through the integration of blue-green infrastructure. Effective biodiversity monitoring is therefore essential to evaluate ecological conditions within these novel socio-ecological systems. Although urban biodiversity monitoring is challenged by its high landscape heterogeneity, dense human populations provide opportunities for large-scale data collection through public participation in citizen science. Using data from 25 City Nature Challenge (CNC) projects across the United Kingdom (2020–2025), we assessed the effects of the four-day bioblitz on species inventories, participation in biological recording, and spatial patterns of recording effort. CNC events doubled public participation in iNaturalist recording relative to baseline activity, leading to the documentation of numerous previously unrecorded species through increased observer effort and broader use of urban blue–green spaces. These results show that CNC events enhance urban biodiversity datasets by increasing the number of observers and reducing spatial and observer biases, providing a cost-effective tool for enriching urban biodiversity data. In addition to generating ecological data, CNC events could have public health benefits through increased exposure to urban blue–green spaces.

## Introduction

Global biodiversity is declining at an unprecedented rate, driven by rapid environmental change and increasing human pressures across all ecosystems (Ceballos et al., 2015; WWF, 2024). At the same time, human populations now exceed 8 billion (United Nations, 2025), and urban development is rapidly expanding, creating novel ecological conditions and conservation challenges (Teixeira & Fernandes, 2020). Understanding how biodiversity is distributed, monitored, and experienced in cities is therefore critical for addressing contemporary ecological questions related to biodiversity loss, ecosystem functioning, and climate resilience, and informing sustainable urban management. In this context, locally grounded knowledge alongside widespread public participation in citizen science offers important opportunities for generating ecological data at the spatial and temporal scales needed (Miller-Rushing et al., 2012; Theobald et al., 2015; Bonney et al., 2023).

Citizen science, defined as scientific research conducted with the participation of non-specialists (OED, 2014), has played a long-standing role in documenting biotic and abiotic environmental change (Miller-Rushing et al., 2012). Such records can span exceptionally long time periods. For example, phenological observations of flowering cherry blossom trees in Japan date back to 801 AD (Primack et al., 2009), while early issues of the Royal Society’s *Journal of Philosophical Transactions*, founded in 1665, were largely based on observational reports from natural philosophers, travellers, and historians documenting phenomena such as weather patterns, animal migrations, and disease outbreaks (Dernham, 1708; Nettleton, 1724; Barr & Saunders, 1777; McDougall-Waters et al., 2015). Prayer records maintained by religious institutions similarly provide long-term environmental observations, which have been used to reconstruct climatic conditions and associated social impacts (Guevara-Murua et al., 2018).

Contemporary citizen science initiatives now generate biodiversity data at unprecedented spatial and temporal scales, spanning diverse taxa and ecosystems. For example, the Audubon Society’s Christmas Bird Count contributes tens of millions of bird records annually (Bateman, 2024), while platforms such as SeagrassSpotter, the Great Eggcase Hunt, and Zooniverse facilitate the global collection of biodiversity data across terrestrial and marine systems (Allen & Gordon, 2023; Seagrass Project, 2025; Zooniverse Team, 2025). Accelerating biodiversity declines and pervasive human influence across ecosystems (Ellis, 2011; WWF, 2024; Holman et al., 2025) necessitate large-scale approaches to biodiversity monitoring, which outpace the capacity of traditional scientific approaches (Pimm et al., 2014). Citizen science helps address this challenge by enabling widespread public participation in data collection across all major biomes, making it a vital tool for contemporary environmental research (Theobald et al., 2015; Mason et al., 2025; Mesaglio et al., 2025). This role is particularly important in urban environments, which are difficult to survey due to their spatial heterogeneity across public, private, built, and open spaces (Cadenasso et al., 2007; Kobori et al., 2016), yet support approximately 46% of the global population (United Nations, 2025). As a result, citizen science data are increasingly central to ecological research, environmental policy, and resource management in urban areas, where high population densities offer potential for the generation of large-scale biodiversity datasets (Ruiz-Gutierrez et al., 2021; Bonney et al., 2023; Mesaglio et al., 2025).

When compared to traditional research methods, citizen science projects can face greater challenges related to uneven sampling effort, variations in data accuracy, and the use of non-standardised equipment (Geldmann et al., 2016; Kosmala et al., 2016). However, technological developments, particularly the internet, have enhanced communication between experts and non-experts, enabled robust systems for data storage and management, and created platforms to disseminate information and coordinate large-scale participation (Dickinson et al., 2012; Kobori et al., 2016). More recently, smartphones with high-resolution cameras, GPS, and integrated apps have become powerful research instruments, enabling users to capture, document, and curate high-quality observations (Teacher et al., 2013). When combined with technological tools, training, and standardised data collection protocols, volunteer surveys can yield results comparable to expert surveys (Holt et al., 2013; Bonney et al., 2014). Advances in statistical methods have further enhanced the value of citizen science data by enabling researchers to leverage large sample sizes while accounting for inherent biases (Kelling et al., 2009). Today, citizen science is widely recognised as a robust approach for investigating species range expansions, phenological shifts, population trends, and invasive species dynamics (Primack et al., 2009; Abdul-Wahab et al., 2024; Pocock et al., 2024) and, to a degree, mortality events and disease ecology (Taylor et al., 2024), offering valuable insights across landscape and macro-ecological scales.

Simultaneously, advances in machine learning and computer vision (artificial intelligence capable of interpreting and analysing images) have improved automated species identification, reducing barriers to participation in citizen science activities and alleviating identification bottlenecks caused by a global shortage of taxonomists (Wäldchen & Mäder, 2018). These innovations have expanded citizen science beyond traditional natural history communities, democratising participation by welcoming individuals with little or no formal training and empowering contributions from people across diverse demographic, educational, and sociocultural backgrounds (Newman et al., 2012). The development of digital tools has enabled the creation of ecological records on an unprecedented scale. For example, citizen science contributions to the Global Biodiversity Information Facility (GBIF) have increased from 11% of records in 2007 to 65% in 2020 (Heberling et al., 2021). Today, the in-kind value of records generated by English-language citizen science projects is estimated at $2.5 billion annually (Theobald et al., 2015).

iNaturalist is one of the most successful citizen science platforms in terms of scale, taxonomic diversity, and global participation (Mesaglio & Callaghan, 2021; Mason et al., 2025). Launched in 2011 as a free mobile application (Seltzer, 2019; Natarajan, 2025a), it became a joint initiative of the California Academy of Sciences and the National Geographic Society in 2017 and was formally registered as a non-profit organisation in 2023 (Natarajan, 2025a). The platform contributes records for 42% of all species represented in GBIF (Loarie, 2023), and, as of 2026, hosts more than 298 million verifiable observations supported by photographic or audio evidence, GPS coordinates, and time-stamped metadata. A central feature of iNaturalist is its continually improving computer vision model, which provides automated species suggestions based on visual traits and geographic context (Loarie, 2025). Guided by these suggestions, users can submit observations and participate in data verification by suggesting or confirming identifications via the app or website. Observations are open access by default and widely used in research, but users can choose to restrict accessibility.

A common format for intensive, place-based biodiversity recording on platforms such as iNaturalist is the bioblitz: a short, time-bounded event that uses crowdsourced observations to document as many species as possible within a defined area. Bioblitzes combine ecological data collection with public engagement, education, and outreach, and are increasingly used in urban environments to raise awareness of local biodiversity while generating large volumes of occurrence data over short time periods (Kobori et al., 2016; Roger & Klistorner, 2016; Tiago et al., 2024). The most prominent demonstration of iNaturalist’s capacity to mobilise public engagement at scale is the City Nature Challenge (CNC) (Palma et al., 2024), an annual four-day global bioblitz in which cities compete to record the greatest number of species and observations, submitted via the iNaturalist platform, within defined boundaries (City Nature Challenge, 2025a). Beginning in 2016 with two projects, it now involves hundreds of cities across six continents, has identified over 70,000 species, and engaged more than 100,000 participants (City Nature Challenge, 2025b). In the United Kingdom (UK), the first CNC took place in 2018 with three cities participating, increasing to 32 cities by 2025. The CNC datasets provide a valuable opportunity to examine how friendly competition can influence user engagement with biodiversity recording and shape ecological datasets. Although framed as a city-based initiative, CNC boundaries have no formal guidelines and often extend beyond urban cores into peri-urban and less-developed landscapes. As a result, the extent to which the event is truly “urban” varies substantially between locations.

Despite the rapid growth of the CNC, limited research has examined how the event contributes to ecological data in urban environments or how patterns of participation shape its scientific and social value (Palma et al., 2024). Engagement-focused bioblitzes like the CNC offer important opportunities to understand how large, semi-structured citizen science events generate ecological knowledge and influence user behaviour. In this study, we use CNC data from the UK to address four questions: (Q1) How urban is the City Nature Challenge? (Q2) To what extent does the CNC contribute new recorded species to the existing iNaturalist dataset? (Q3) To what extent does the CNC increase user engagement on iNaturalist? and (Q4) Does participation in the CNC promote increased visitation and biodiversity recording within urban blue–green spaces? Together, these questions clarify the ecological value of the event and demonstrate how short-term shifts in recording behaviour can shape patterns of public engagement with urban biodiversity.

## 2. Methods

### 2.1 City selection and urban land cover assessment

CNC projects within the UK active between 1 October 2020 and 1 October 2025 were identified using iNaturalist leaderboard summaries (see Supplementary Table 1). Projects were retained for analysis if they had participated in at least two CNC events during this period. Boundaries were defined using unique numeric project codes and downloaded directly from project-specific URLs (see Supplementary Table 2). Urban areas associated with each project were quantified using the Office for National Statistics Built Up Extents layer (Office for National Statistics, 2024). Built-up area polygons falling within each CNC project area were extracted using a cookie-cutter approach in QGIS v3.36.3 (*vector-geoprocessingtools-clip*). The proportion of urban cover was then calculated for each project (*field calculator-$area*). Greater Belfast was excluded from the urban assessment due to missing official built-up area data.

### 2.2 iNaturalist data preparation

All observations classified as *Verifiable*— i.e. records of wild organisms which are supported by photographs or sound recordings, along with geographic coordinates and date-time metadata—were retrieved for each project using iNaturalist’s Explore and Export functions. For projects exceeding 200,000 observations, data were exported in multiple batches due to download limits. Only records with the following licenses were retained for analysis: CC0, CC-BY, CC-BY-NC, CC-BY-SA, CC-BY-ND, CC-BY-NC-ND, CC-BY-NC-ND. Records with blank license fields were excluded.

For all analyses requiring species-level resolution, observations were filtered to iNaturalist’s Research Grade classification, retaining only records with at least two agreeing user identifications and for which at least two-thirds of all identifications were in agreement. Subspecies designations were reduced to the species level to avoid inconsistencies arising from the variable use of subspecies classifications amongst users (e.g., *Passer domesticus domesticus* to *Passer domesticus*). All hybrid taxa were retained. For analyses of user participation and engagement, all verifiable records were included, acting as proxies of activity. In spatial analyses, only records with a location accuracy of 20 metres or less were retained.

### 2.3 Species Accumulation and Difference Curves

To evaluate the contribution of CNC events to iNaturalist-documented taxonomic richness, species accumulation curves (SACs) were generated for each project by aggregating observations into monthly intervals and calculating the cumulative number of unique species through time. For each city, two SACs were produced: one that included all observations, and another that excluded observations from CNC events. Difference curves were then calculated from the two SACs. Finally, the annual difference in cumulative species between datasets with and without CNC observations was compared across successive years of CNC participation. When projects had started before the 2020-2025 window, previous years were accounted for in the analysis (i.e., for projects which started in 2018, 2020 would be classed as year three in our analysis).

### 2.4 Unique CNC Taxa

Taxonomic summaries of species exclusively recorded in each CNC event were compiled using iNaturalist’s iconic_taxon_names. Records with missing or indeterminate classifications were removed, and entries labelled “Animalia” were reassigned to “Other Animalia” to distinguish them from specific animal groups. For interpretation, taxon common names were adopted from original classifications (see Supplementary Table 3).The effects of project size, percentage of urban cover, and relative CNC user increase on the number of unique taxa were also explored using statistical models (see 2.8 Data Analysis section).

### 2.5 User Engagement

User participation and behaviour were assessed using a weekly sliding window spanning five weeks before and five weeks after each CNC event, with the event itself defined as week 0. Week 0 encompassed the four-day CNC period (Friday–Monday) as well as the three preceding days (Tuesday–Thursday) to capture pre-event activity associated with practice observations, app familiarisation, and organised training sessions. Using this structure, we quantified engagement as the number of unique users contributing observations each week. Baseline activity was defined as the mean weekly number of unique users during the five pre-event weeks (weeks −5 to −1). CNC engagement was then calculated as a proportion of this baseline.

### 2.6 City Case Study: Hull, UK

To explore the combined effects of user behaviour and land cover, spatial analyses were conducted for the Hull CNC project, a predominantly urban project that has participated in every CNC since 2023. Spatial patterns of recording activity were assessed using 250 m grid cells over the same sliding-window period described above. Accessibility of inactive grids was assessed using the Ordnance Survey MasterMap Highways Network; grid cells that did not overlap with public roads or footpaths were manually excluded from analysis, as these areas were considered inaccessible to participants. Each grid was classified as *Active on iNaturalist* (used both before and during the event), *CNC-only* (used exclusively during the CNC), or *inactive* (accessible but never used during or outside of the event). All active grid categories (*Active on iNaturalist* and *CNC-only*) contained records from at least one user, whereas *inactive* grids contained no observations.

Land cover for each grid was quantified from a composite single-band raster layer (1 m resolution) created using high-resolution land-use data (Ordnance Survey, 2023), the European Space Agency *WorldCover* map 2021 (10 m resolution; Zanaga *et al*., 2022), and manual digitisation.

First, Ordnance Survey (OS) land-use vector layers were used to delineate Buildings, Gardens, Roads (line features buffered by 5 m), Amenity Grassland, Surface Water, Tidal Water, Foreshore, and Crops. Remaining gaps were then filled using *WorldCover* data, which provided coverage for Trees, Impervious Surfaces, and Grasslands (where absent), without replacing OS layers. Manual digitisation, informed by Google Satellite imagery, local field-based knowledge, and existing habitat records, was used to add Rough Grassland, Riparian Vegetation, and Saltmarsh. The resulting composite was rasterised to a 1 m resolution and classified into 15 land-cover categories. For broader analyses, these categories were aggregated into green, grey, and blue cover (see Supplementary Table 4).

### 2.7 Data Analysis

All analyses were conducted in R (version 4.3.3; R Core Team, 2025) and the specified packages. Species accumulation analyses were conducted following the collector’s principle, under which cumulative species richness is expected to approach an asymptote with increasing sampling effort (Colwell et al., 2004). Difference curves were calculated as the cumulative difference in species richness between SACs with and without CNC observations for each city. A linear mixed-effects model (LMM) from the *lme4* package (Bates et al., 2015) was used to quantify these effects over successive years. Summary statistics were calculated to describe the composition of taxa recorded exclusively during CNC events. A negative-binomial generalised linear model (nb.GLM) from the MASS package (Venables & Ripley, 2002) was used to examine variation in the number of unique taxa across projects, using the CNC–baseline difference in users, project area (km²), and urban cover (%) as predictors. Multicollinearity among predictors was assessed using variance inflation factors (VIFs) and pairwise correlations (see Supplementary Figure 1). Model assumptions for the LMM and nb-GLM were checked using simulation-based residual diagnostics from the DHARMa R package (Hartig, 2024).

Temporal patterns of user participation were modelled using linear regression, with weeks relative to the CNC event included as a continuous predictor and a quadratic term to capture non-linear trends in engagement. CNC engagement was expressed relative to the pre-event baseline, and week 0 (the CNC week) was included as a categorical indicator to explore the effect of the event on participation. Model performance was evaluated through visual inspection of residuals and comparison of fitted and observed values.

Spatial analyses were conducted at the 250 m grid-cell level using the R package *sf* (Pebesma & E, 2023). Grid classifications (*active on iNaturalist, CNC-only*, *inactive*) and land-cover proportions (*green*, *grey*, *blue*) were compared to assess relationships between user activity, species richness, and the local environment. Land-cover diversity was also estimated for each grid cell based on 15 land-cover classes (Supplementary Table 4), using the Shannon’s diversity index in the R package vegan (Oksanen et al., 2025). Land cover across the three grid categories was summarised using descriptive statistics, and grid-level differences were assessed with non-parametric bootstrapping (1,000 replicates) implemented in R using the boot package (Canty & Ripley, 2025). To account for unequal sampling effort across grid categories, bootstrapping was standardised to the smallest group size (141 CNC-only grids), with an equal number of grids randomly resampled from each category during each iteration.

## 3. Results

### 3.1 UK-Level Analysis

The combined UK dataset contained 3,080,822 observations (mean per project = 123,232; range = 9,948–604,072), contributed by 100,508 observers (mean per project = 4,020; range = 358–20,832). Observations removed due to blank licences accounted for 15.1% of records. CNC participation histories ranged from two to eight years at the project-level (Supplementary Table 5).

#### 3.1.1 Extent of Urban Cover Across CNC Projects

In total, 25 projects participated in at least two CNC events within the five-year comparison window (2020–2025). Project areas differed widely, ranging from 62 to 10,510 km². Only six of the 25 projects had over 50% urban cover. The degree of urbanisation was highly variable, averaging 26.5% and ranging from 6 to 75%. Nottingham, Birmingham, Hull and Coventry ranked highest in urban cover, whereas Suffolk, North East England and Glasgow were among the least urban (Fig. 1).

**Fig. 1:**
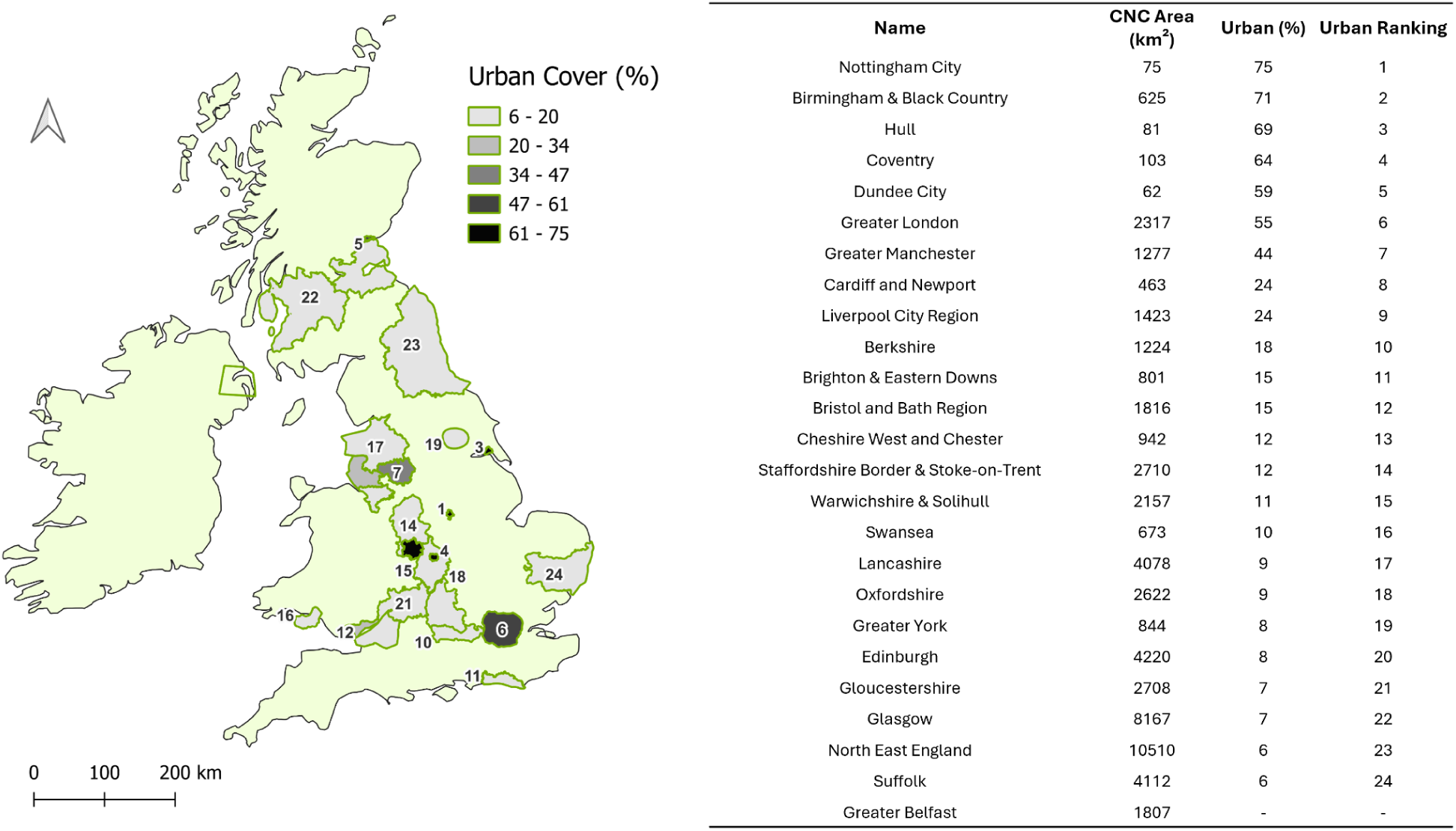
Left: Map of the UK showing CNC project boundaries coloured by their percentage of built (urban) land cover. Right: Table summarising CNC project area (km²), built cover (%), and urban ranking. Projects are listed by decreasing proportion of urban cover.

#### 3.1.2 Contribution to Existing Biodiversity Records

Species accumulation curves confirmed that CNC events contributed unique ecological records across all projects, at varying scales (Supplementary Figure 2). Difference curves further revealed that peaks in CNC-associated gains in cumulative species richness slowly declined between events and decreased in magnitude across successive years (see Fig. 2a; Supplementary Figure 3). The long-term pattern indicates an average decline of approximately 0.98% in unique species richness per year of participation (LMM β = −0.98 ± 0.24 SE, *t* = −4.08, *p* <0.001) (see Fig. 2b). Simulation-based residual diagnostics (DHARMa) showed no evidence of model misfit (see Supplementary Figure 4 and Supplementary Table 6 for model summary). A nonlinear model was used for comparison, but it did not improve model fit and the quadratic term was not significant. Mean CNC-related gains were highest in the first three years of participation (∼5–6%) and declined in subsequent years, reaching around 2–3% by years 5–8 (see Supplementary Table 7).

**Fig 2:**
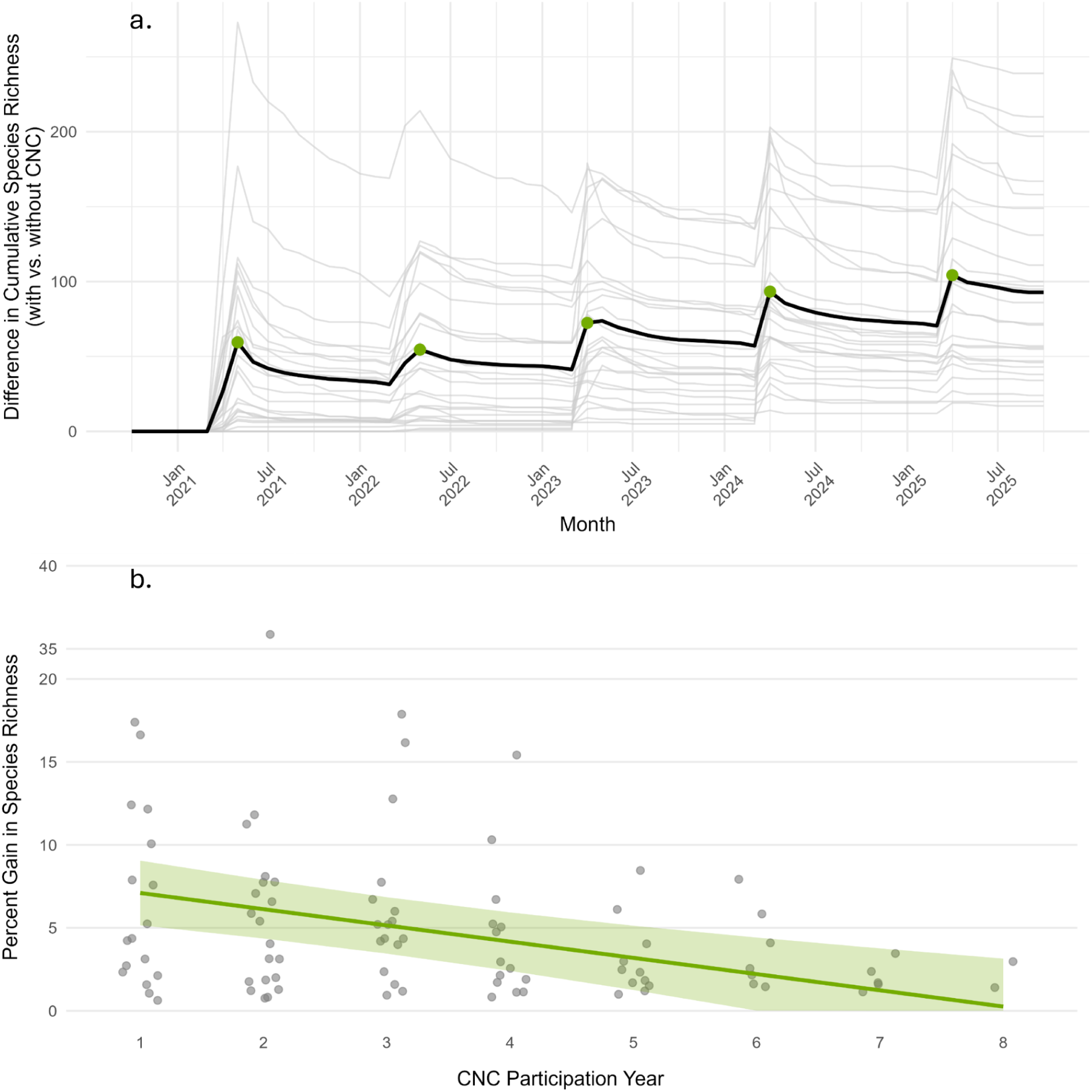
a. Difference in cumulative species richness (with vs. without CNC data) across the five-year study period for each individual CNC project (pale grey lines) (n = 25). The bold black line shows the monthly mean across all projects active during that month. Green dots mark CNC events. b. Species richness gains in the iNaturalist dataset (%) attributable to CNC participation across successive years (1–8) for all 25 CNC projects. Grey points represent individual project-year CNC observations (n = 93). The green line shows the fitted LMM trend, and the shaded band indicates the 95% confidence interval for the population-level mean. The model revealed a significant decline in richness gains with repeated participation.

The number of species recorded exclusively during CNC events varied widely among cities (mean = 112 ± 53 SD, range = 34–205; Fig. 3a). Hull, Coventry and York recorded the highest numbers of CNC-unique species, whereas Oxfordshire and Edinburgh recorded the fewest. The only significant predictor of unique taxa found with the nb-GLM was relative user engagement (Estimate: 1.25, SE: 2.31, z value: 4.9, p-value: <0.001; Fig. 3c). Simulation-based residual diagnostics showed no evidence of model misfit (Supplementary Figure 6 and Supplementary Table 8 for summary). On average, CNC-unique records were dominated by *Plants* (mean = 46 %, range = 38–52 %), *Insects* (mean = 34 %, range = 28–42 %), and *Fungi* (mean = 15 %, range = 9–21 %; Fig. 3b), with smaller contributions from other groups. Full city-level values are provided in Supplementary Table 9.

**Fig. 3:**
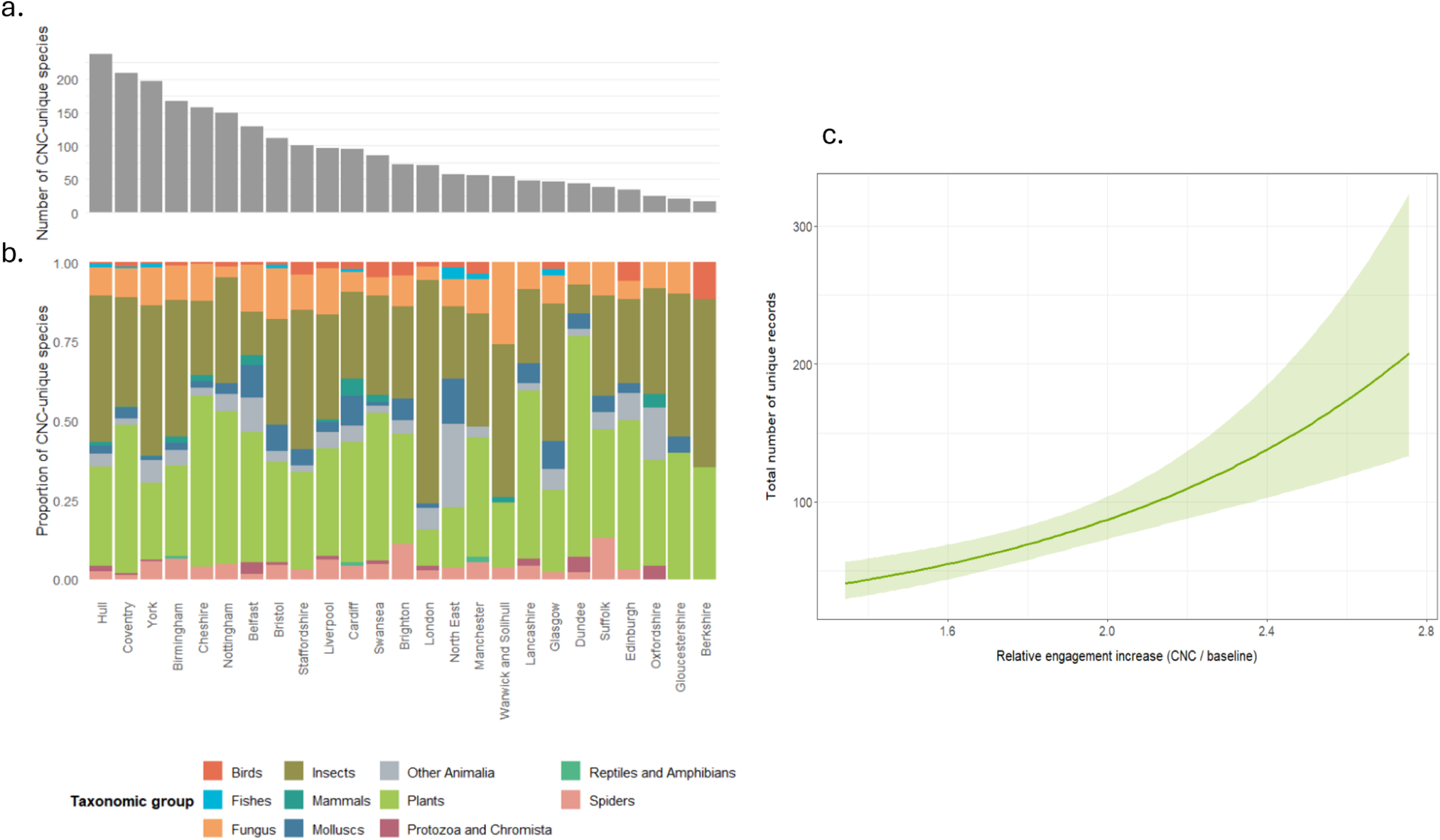
a. Total number of CNC-unique species recorded in each participating city, in descending order. b. Composition of major taxonomic groups contributing to CNC-unique biodiversity within each project. c. Effect of relative user engagement extracted from the nb-GLM (Estimate: 1.25, SE: 2.31, z value: 4.9, p-value: <0.001).

#### 3.1.3 Effect of the CNC on User Engagement

Across the UK, mean user engagement during the CNC was approximately 90% higher than the pre-event baseline (range = 34–176%). This pattern was strongly supported by the linear model (β = 96.5 ± 6.0, R² = 0.71, p < 0.001), see Fig. 4. Model assumptions were assessed using standard residual diagnostics and showed no major violations (see Supplementary Figure 7 and Supplementary Table 10 for model details).

**Fig. 4:**
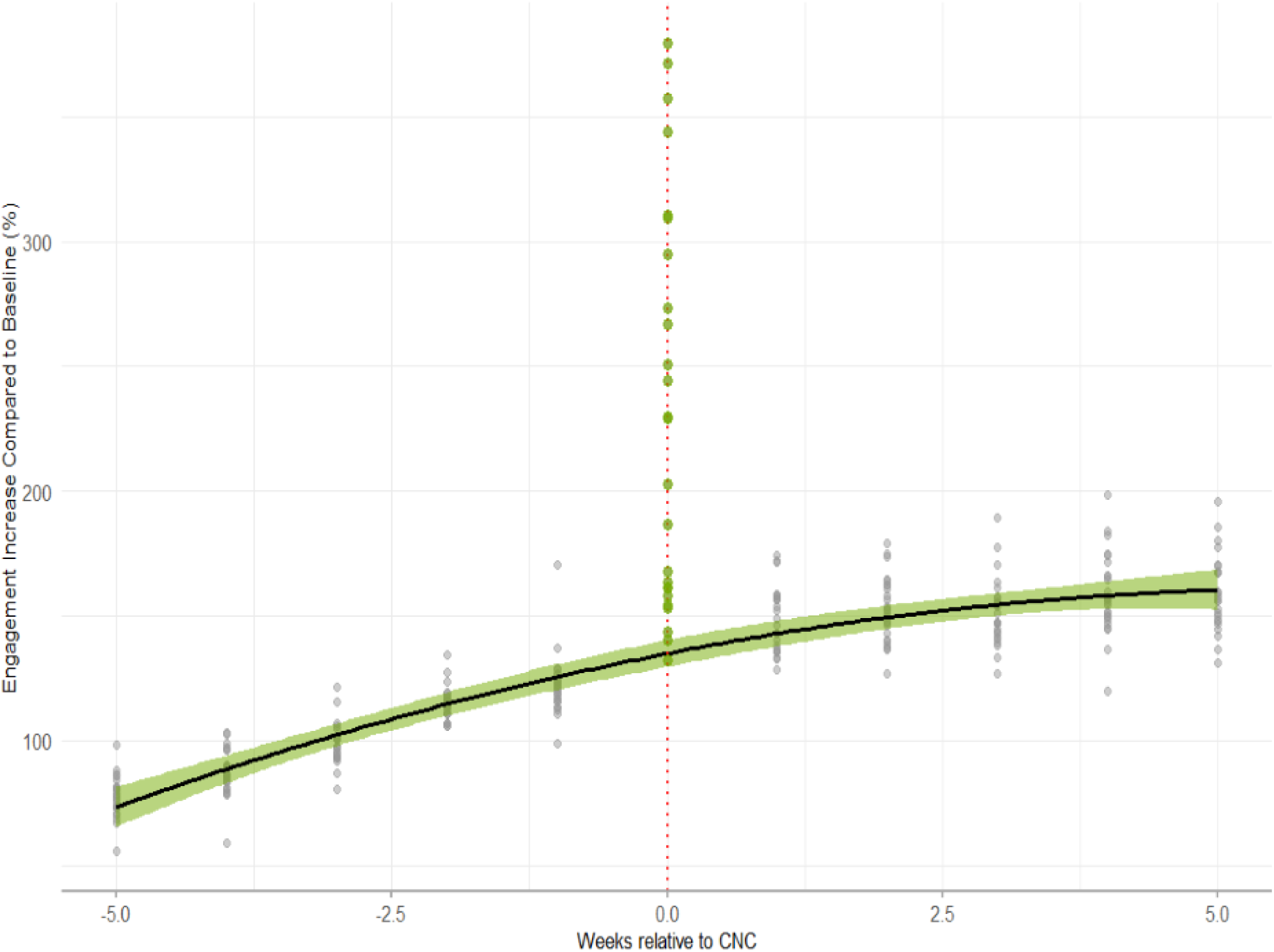
Weekly iNaturalist user activity before (−), during (0), and after the CNC week, relative to the seasonal baseline (Y-axis). Grey points show weekly user engagement, and green points highlight engagement during CNC week. Black curve shows the fitted quadratic regression with week 0 removed.

### 3.2 User Behaviour During the CNC: A Case Study of Hull

Spatial analysis of 1,016 accessible grid cells within Hull identified 141 cells used exclusively during CNC events (14%), 411 active both before and during the CNC (40%), and 464 that remained inactive throughout the study period (46%) (Figure 4). Land-cover summaries showed that CNC-only grids had the highest proportion of blue cover, followed by baseline grids, while inactive areas contained the least. Green cover dominated all grid types,followed by grey surfaces. See Table 2.

**Table 2:**
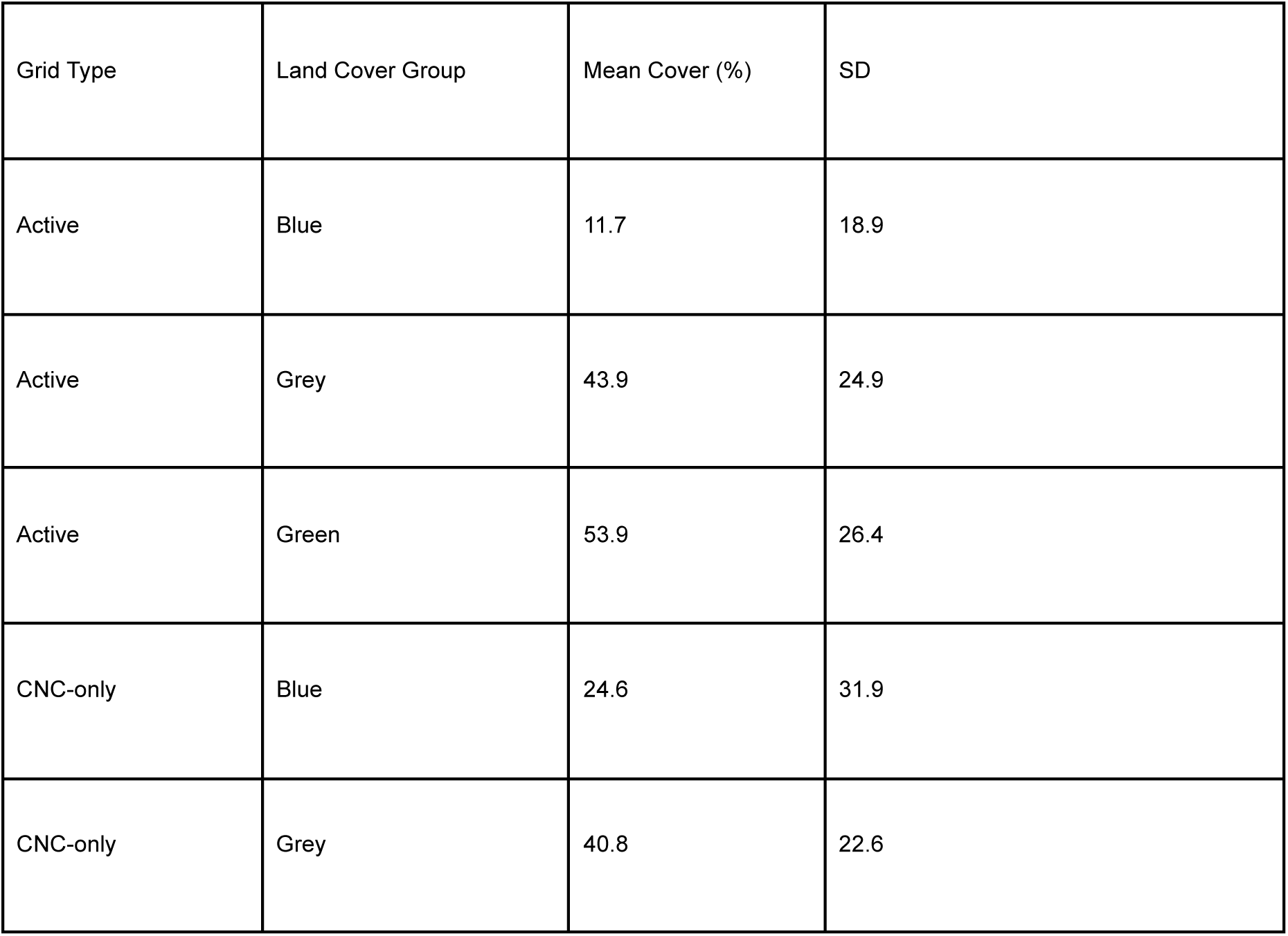
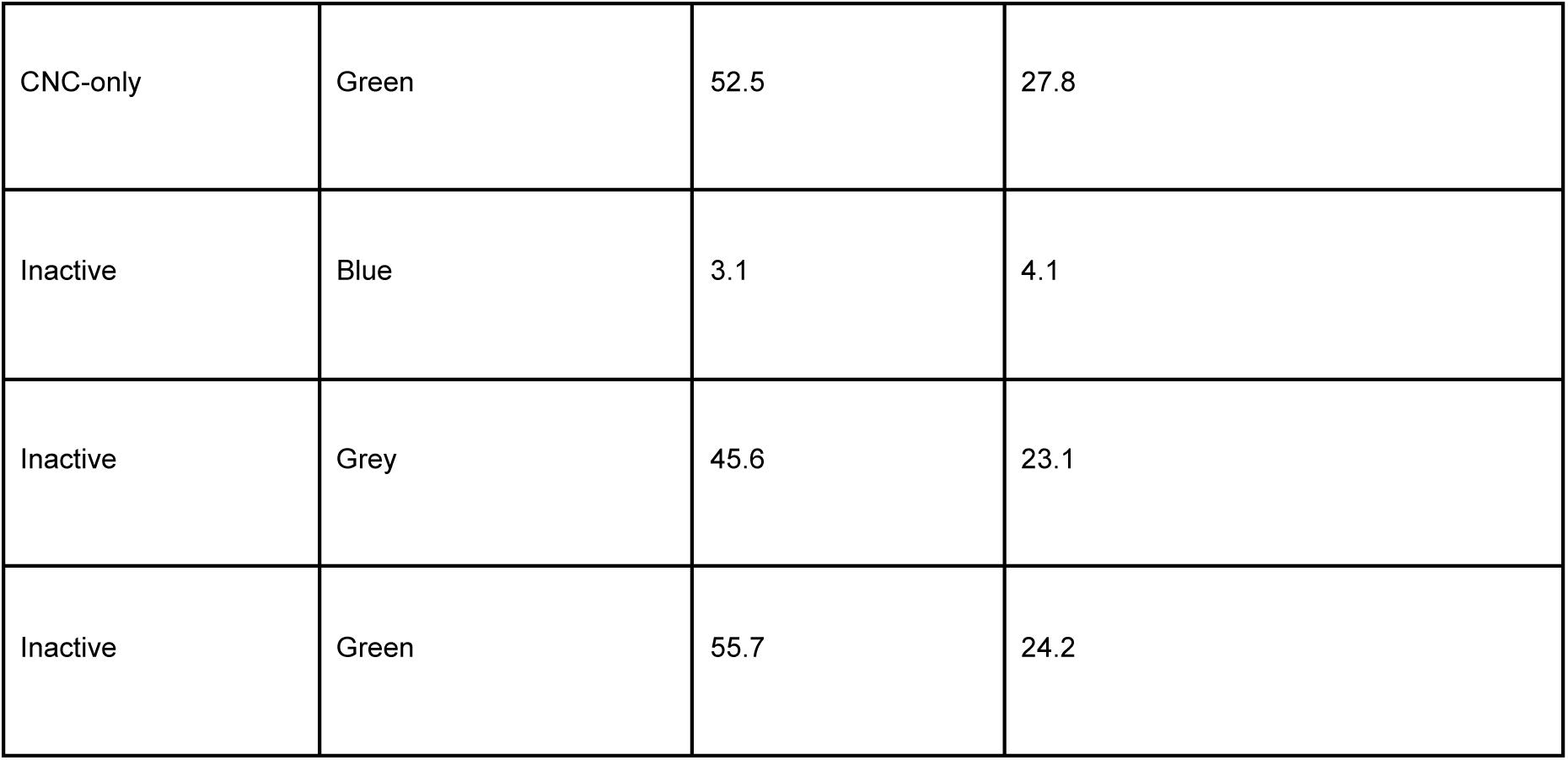
Summary of land cover groups by grid type.

| Grid Type | Land Cover Group | Mean Cover (%) | SD |
| --- | --- | --- | --- |
| Active | Blue | 11.7 | 18.9 |
| Active | Grey | 43.9 | 24.9 |
| Active | Green | 53.9 | 26.4 |
| CNC-only | Blue | 24.6 | 31.9 |
| CNC-only | Grey | 40.8 | 22.6 |
| CNC-only | Green | 52.5 | 27.8 |
| Inactive | Blue | 3.1 | 4.1 |
| Inactive | Grey | 45.6 | 23.1 |
| Inactive | Green | 55.7 | 24.2 |

Bootstrapping revealed that CNC-only grids had significantly higher proportions of blue cover compared with both inactive grids (median difference = 0.215, 95% CI = 0.135–0.297) and shared grids (median difference = 0.124, 95% CI = 0.0287–0.228; Fig. 5). However, CNC-only grids were significantly less diverse in land cover (Shannon’s index) than active grids (median difference = −0.110, 95% CI = −0.197−0.033). All other bootstrap comparisons were not statistically significant. See Figure 5 and Supplementary Table 11.

**Fig. 5:**
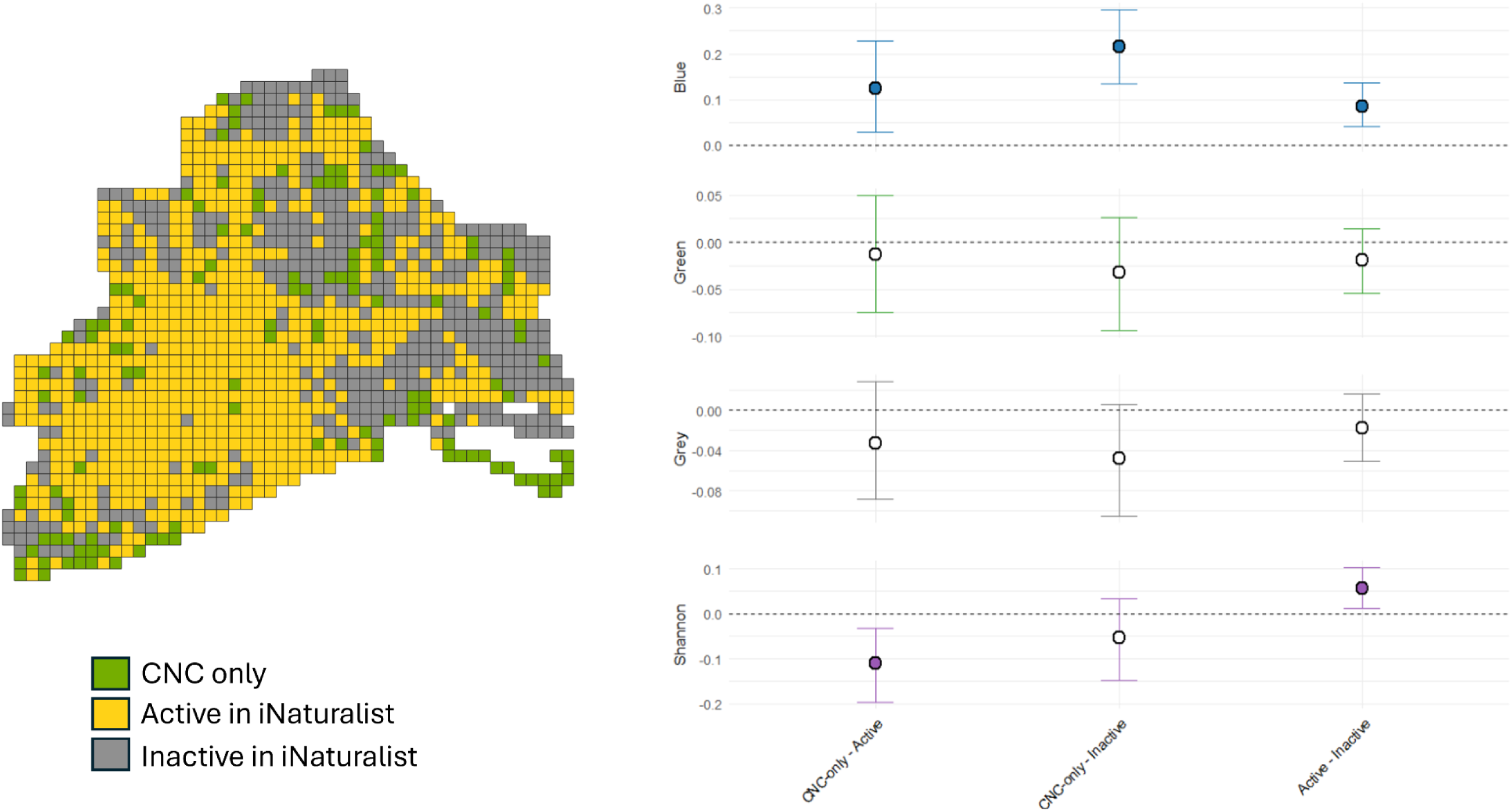
Left: Hull map showing 250 m grid cells classified according to iNaturalist activity status. Right: Bootstrapped mean differences (± 95% confidence intervals) in blue, green, grey land cover and Shannon’s diversity for each grid-cell category. Statistically significant differences are shown as filled points

## 4. Discussion

In the face of global biodiversity decline, it is critical that urban development integrates biodiversity into sustainable design, and that these interventions are monitored to assess their effectiveness. To achieve this, ecological data from these complex understudied systems are required. We show that the City Nature Challenge (CNC) delivers meaningful ecological benefits, generating rapid increases in biodiversity records and revealing species that would otherwise remain undetected. In addition, the CNC appears to influence participant behaviour by encouraging exploration and recording of biodiversity within urban blue–green spaces. These behavioural responses expand both the spatial and taxonomic coverage of urban biodiversity data, while simultaneously increasing opportunities for direct contact with nature. Collectively, these outcomes highlight the CNC’s capacity to generate ecologically valuable data while shaping how people engage with urban environments.

### 4.1 CNC Urban Extent

We found that CNC project boundaries in the UK often encompass extensive areas of undeveloped land, including peri-urban green spaces, agricultural landscapes, and protected areas, despite the initiative’s focus on cities. This likely reflects efforts by CNC organisers to maximise participation and to include areas designated for their biodiversity value, such as nature reserves. For example, the York CNC boundary encompasses several large Yorkshire Wildlife Trust nature reserves more than 10 km from the city itself, while the North East of England project covers a vast area including Northumberland National Park, extensive coastline, and rural landscapes. Although this does not pose a major issue for urban researchers, who can filter urban records retrospectively, it likely draws potential observer effort away from cities. This is notable, given that cities are already understudied when compared to natural landscapes, particularly protected areas (Miller & Hobbs, 2002; Martin et al., 2012). Expanding project ranges beyond city boundaries, therefore, risks weakening the CNC’s usefulness for understanding and monitoring urban biodiversity. In this study, we used original project boundaries to show that the degree of urbanisation varies substantially across UK projects. As a result, comparisons of CNC dataset contributions primarily reflect the influence of the CNC rather than isolating urban-specific effects. Future work aiming to disentangle urban impacts may therefore benefit from refining project selections to those with high urban cover or extracting data from within city boundaries.

### 4.2 Contribution to Biodiversity Knowledge

Although each CNC year contributes unique species (previously unrecorded in the area in iNaturalist), the extent of CNC’s contribution of unique species diminishes over successive years, with most additions occurring in the first years. This pattern is consistent with the well-known saturation of species accumulation in biological recording (Colwell et al., 2004; Kelling et al., 2015) and suggests that projects with a strong baseline of iNaturalist recorders are less likely to find unique species than projects with lower baseline engagement. This was shown in projects that saw a substantial surge in participation during the event, such as Hull and Coventry. This pattern highlights the interplay between existing recording effort and the incremental value of short, intensive bioblitz-style events. As iNaturalist Research Grade records are sent to the Global Biodiversity Information Facility, these new records also have broader conservation value.

The taxonomic composition of CNC-unique taxa was broadly consistent across projects, despite large differences in project size and urbanisation. Plants and insects contributed the largest proportions of these unique records, suggesting that species richness within these groups remains far from saturated across locations. Plants and insects are also the most recorded taxa on iNaturalist (Natarajan, 2025b). However, both groups remain underrepresented on the platform relative to their known species diversity; many plant and insect classes still show low coverage overall, in contrast to birds, for which more than 80% of the world’s ∼11,000 species have been documented on the platform (Di Cecco et al., 2021). Their prominence in CNC datasets reflects their diversity, and suitability for detection and identification using mobile-phone photography (Di Cecco et al., 2021), the primary recording tool for most participants. Fungi and arachnids also contributed notable numbers of unique taxa, whereas iconic groups such as mammals accounted for comparatively few. Considering that citizen science datasets often exhibit biases toward large, conspicuous, or easily identifiable species (Callaghan et al., 2021), the CNC appears to counterbalance some of these tendencies by capturing a broader range of typically under-recorded organisms, thereby helping to reduce taxonomic sampling biases.

### 4.3 User engagement and behaviour

The substantial rise in the number of people recording during the CNC period (+ 98%) confirms that the event itself drives a marked increase in participation beyond what would be expected from seasonal trends in user activity (Di Cecco et al., 2021). This indicates that the CNC is an effective tool for increasing short-term engagement. However, the highly concentrated nature of this activity also suggests that increased user engagement is mostly temporary, a pattern seen in other citizen science programmes (Eveleigh et al., 2014). Future work could examine whether such events lead to sustained changes in user behaviour, particularly among participants who return for multiple annual CNC events, and whether any long-term effects accompany the periodic surges generated during the event.

We found that the majority of CNC observations in the Hull CNC project come from locations already active in the iNaturalist recording community, likely reflecting sites which have good access, known biodiversity value or an established reputation as productive places to record (Callaghan, Poore, et al., 2019; Callaghan, Rowley, et al., 2019). However, some locations were only active during CNC dates, suggesting that the event provides opportunities, motivation or access to areas that are otherwise under-recorded outside of CNC events.

Patterns of land-cover use across locations active in iNaturalist recording highlight a strong preference for blue-space environments. This was most evident in locations only active during the CNC, which contained substantially more blue-space cover than inactive locations. These results indicate that participants are drawn to record in urban blue spaces such as rivers, drains and ponds, reflecting their awareness of the diverse and distinctive species assemblages in these areas (Morgan et al., 2025). In contrast, overall land-cover diversity was lower in CNC-only locations compared with grids active during routine iNaturalist recording. This suggests that although habitat diversity shapes routine recorder behaviour, the CNC’s time-limited structure focuses effort on under-recorded blue-space environments, which can be more homogeneous than other urban habitats.

Importantly, these behavioural shifts have implications beyond biodiversity data collection. Blue space exposure has strong links to improved health and well-being (White et al., 2020; Brückner et al., 2021; Wang et al., 2022); therefore, participation in the CNC may provide incidental benefits by increasing users’ contact with these restorative environments. Similarly, benefits have been observed in citizen-science recording more broadly, where purposeful engagement with nature can enhance psychological and physical well-being (Hobbs & White, 2012; Domroese & Johnson, 2017). Thus, the CNC likely supports both biodiversity engagement and opportunities for nature contact in urban areas, providing associated human-health benefits and underscoring its wider social value.

### 4.4 Practical Implications

Both the ecological data and participant behavioural patterns observed in the CNC are highly relevant to local planners, conservation practitioners and government agencies seeking to promote public engagement with nature and deliver equitable, sustainable cities. In England, national environmental policy initiatives such as Local Nature Recovery Strategies and Biodiversity Net Gain, both key delivery mechanisms under the Environmental Improvement Plan (UKGOV, 2025), are placing increasing emphasis on urban biodiversity. Gaining insight into how people use and experience nature in cities is increasingly recognised as essential for meeting statutory requirements and strategic targets, reflecting recent rewilding policy discussions that position inclusive public engagement and citizen-generated evidence as integral to effective planning and policy implementation (Hopkins et al., 2026).

Sustainability agendas and urban research increasingly discuss blue–green infrastructure and nature-based solutions in response to climate and biodiversity pressures (Kabisch et al., 2016, 2017; United Nations, 2022). However, Frantzeskaki et al. (2019) highlight that building a robust evidence base to evaluate the effectiveness of nature-based solutions remains a challenge, particularly in assessing their performance across contexts, trade-offs, and long-term impacts. Part of this challenge lies in monitoring ecological conditions at an appropriate resolution and scale across complex urban systems (Grimm et al., 2008; Kabisch et al., 2016). In this context, citizen science can play an important role, with initiatives such as the CNC contributing spatially extensive species occurrence datasets at high spatial resolution, supporting the evaluation and adaptive management of blue–green strategies over time.

From a local planning perspective, where resources are often limited, the CNC offers a cost-effective way to increase biodiversity records at scale. The benefits are most evident in the first years of participation, which generate substantial influxes of new ecological records, indicating that even short-term involvement can provide meaningful returns. Given the scale and complexity of urban environments, few approaches outside citizen science can produce datasets of comparable scope (Theobald et al., 2015), and the CNC delivers an intensified version of the value already demonstrated by year-round citizen-science platforms (Mason et al., 2025; Mesaglio et al., 2025).

The CNC also provides insight into where people choose to record biodiversity, revealing underused green and blue spaces and accessibility issues relevant to urban management strategies. These spatial patterns also reflect public perceptions of where biodiversity is most likely to be encountered within urban landscapes. In our study, recording activity was strongly associated with blue-space environments, aligning with evidence of frequent encounters with urban wildlife and public associations between water and biodiversity (Chapter 4). This points to opportunities for integrating water features, bio-SUDS (Monberg et al., 2018), blue-space regeneration (Brückner et al., 2021), or establishing “blue–green” ecological corridors within urban design (Hyseni et al., 2021; Donati et al., 2022).

These behavioural patterns are also relevant to the growing field of green social prescribing, which encourages nature-based activities to support physical and mental health (Bragg & Leck, 2017; Robinson et al., 2020). Participation in the CNC provides a semi-structured, accessible pathway for individuals to engage with natural environments, particularly restorative urban green and blue spaces, while also encouraging shared participation and social interaction, both of which have been associated with benefits for health and wellbeing (Jennings & Bamkole, 2019; White et al., 2019; Wan et al., 2021). CNC outcomes may therefore help identify urban areas suitable for health-oriented interventions, serving as a proxy for accessibility, perceived safety, and user comfort, in addition to ecological quality. This suggests that biodiversity monitoring initiatives and public health strategies offer mutual benefits, building both ecological understanding and social connection in urban environments.

Lastly, participation in biodiversity recording has been shown to strengthen local environmental stewardship and to foster broader environmental awareness and pro-environmental attitudes beyond the immediate study area (Cooper et al., 2007; Hobbs & White, 2012; Santori et al., 2021). These broader benefits are likely to build support for improved care of public blue–green spaces, increased support for rewilding, and greater public backing for green-infrastructure initiatives. Given that participation in iNaturalist approximately doubles during the City Nature Challenge (CNC), short-term, high-visibility events that engage the public beyond the routine user community may provide valuable opportunities to reach new audiences and encourage initial connections with nature.

## 5. Conclusion

Our findings show that the CNC delivers a combination of ecological outcomes and public behavioural responses that can make valuable contributions to urban biodiversity research and environmental policy. Alongside rapid increases in species records, the CNC influences user behaviour by prompting exploration of new locations, particularly blue-space environments, and by increasing nature engagement among new audiences. These behavioural shifts not only enrich ecological datasets but also offer opportunities to enhance environmental stewardship and improve public health through greater nature contact. Within the UK’s emerging environmental policy landscape, the CNC represents a scalable and cost-effective mechanism for improving urban biodiversity knowledge while engaging diverse communities with their local environments. Strengthening the integration of such initiatives into planning, monitoring, and community-engagement strategies could further enhance both their scientific and social value, while also supporting nature-based public health initiatives through increased exposure to urban green and blue spaces.

## Supporting information

See supplementary material here.

## Acknowledgements

We would like to thank the University of Hull for funding this research.

## Data Availability

Code and data has been made available here: https://github.com/MCMorgan06/Blue_Space_CNC_Data

