## Supplementary material for "The City Nature Challenge increases urban biodiversity knowledge and public engagement with blue spaces": See supplementary material here.

**Supplementary Table 1:** iNaturalist leaderboard links.

| | |
|---|---|
| 2021 | https://uk.inaturalist.org/projects/city-nature-challenge-2021-uk-participants |
| 2022 | https://uk.inaturalist.org/projects/city-nature-challenge-2022-uk-leaderboard |
| 2023 | https://uk.inaturalist.org/projects/city-nature-challenge-2023-uk-leaderboard |
| 2024 | https://uk.inaturalist.org/projects/city-nature-challenge-2024-uk-leaderboard |
| 2025 | https://uk.inaturalist.org/projects/city-nature-challenge-2025-uk-leaderboard-umbrella-project |

**Supplementary Table 2:** Download links for all geographic boundaries used in the study.

| **Name** | **Code** | **Download link** |
|---|---|---|
| Birmingham & Black Country Border | 144101 | https://www.inaturalist.org/places/geometry/144101.kml |
| Hull City Council Border | 186496 | https://www.inaturalist.org/places/geometry/186496.kml |

| | | |
|---|---|---|
| Greater London Border | 66357 | https://www.inaturalist.org/places/geometry/66357.kml |
| Liverpool City Region Border | 130732 | https://www.inaturalist.org/places/geometry/130732.kml |
| Bristol and Bath Region Border | 123650 | https://www.inaturalist.org/places/geometry/123650.kml |
| Brighton & Eastern Downs Border | 164862 | https://www.inaturalist.org/places/geometry/164862.kml |
| Staffordshire Border & Stoke-on-Trent | 211134 | https://www.inaturalist.org/places/geometry/211134.kml |
| Cheshire West and Chester Border | 187347 | https://www.inaturalist.org/places/geometry/187347.kml |
| Greater York Border | 197922 | https://www.inaturalist.org/places/geometry/197922.kml |
| North East England Border | 131090 | https://www.inaturalist.org/places/geometry/131090.kml |
| Lancashire | 138194 | https://www.inaturalist.org/places/geometry/138194.kml |
| Greater Manchester | 138285 | https://www.inaturalist.org/places/geometry/138285.kml |
| Greater Edinburgh | 30456 | https://www.inaturalist.org/places/geometry/30456.kml |
| Greater Edinburgh | 30461 | https://www.inaturalist.org/places/geometry/30461.kml |
| Greater Edinburgh | 30463 | https://www.inaturalist.org/places/geometry/30463.kml |
| Greater Edinburgh | 30465 | https://www.inaturalist.org/places/geometry/30465.kml |
| Greater Edinburgh | 30468 | https://www.inaturalist.org/places/geometry/30468.kml |

| | | |
|---|---|---|
| Greater Edinburgh | 30481 | https://www.inaturalist.org/places/geometry/30481.kml |
| Oxfordshire | 30388 | https://www.inaturalist.org/places/geometry/30388.kml |
| Warwickshire, Solihull | 186686 | https://www.inaturalist.org/places/geometry/186686.kml |
| Warwickshire, Solihull | 186687 | https://www.inaturalist.org/places/geometry/186687.kml |
| Sufflok | 30407 | https://www.inaturalist.org/places/geometry/30407.kml |
| Greater Belfast | 198158 | https://www.inaturalist.org/places/geometry/198158.kml |
| Berkshire | 30318 | https://www.inaturalist.org/places/geometry/30318.kml |
| Swansea | 30500 | https://www.inaturalist.org/places/geometry/30500.kml |
| Dundee | 192764 | https://www.inaturalist.org/places/geometry/192764.kml |
| Gloucestershire | 30344 | https://www.inaturalist.org/places/geometry/30344.kml |
| Coventry | 164957 | https://www.inaturalist.org/places/geometry/164957.kml |
| Coventry | 164860 | https://www.inaturalist.org/places/geometry/164860.kml |
| Coventry | 164890 | https://www.inaturalist.org/places/geometry/164890.kml |
| Cardiff and Newport | 163986 | https://www.inaturalist.org/places/geometry/163986.kml |
| Nottingham City | 144218 | https://www.inaturalist.org/places/geometry/144218.kml |
| Greater Glasgow | 30459 | https://www.inaturalist.org/places/geometry/30459.kml |

| | | |
|---|---|---|
| Greater Glasgow | 30460 | https://www.inaturalist.org/places/geometry/30460.kml |
| Greater Glasgow | 30462 | https://www.inaturalist.org/places/geometry/30462.kml |
| Greater Glasgow | 30466 | https://www.inaturalist.org/places/geometry/30466.kml |
| Greater Glasgow | 30467 | https://www.inaturalist.org/places/geometry/30467.kml |
| Greater Glasgow | 30470 | https://www.inaturalist.org/places/geometry/30470.kml |
| Greater Glasgow | 30471 | https://www.inaturalist.org/places/geometry/30471.kml |
| Greater Glasgow | 30474 | https://www.inaturalist.org/places/geometry/30474.kml |
| Greater Glasgow | 30477 | https://www.inaturalist.org/places/geometry/30477.kml |
| Greater Glasgow | 30478 | https://www.inaturalist.org/places/geometry/30478.kml |
| Greater Glasgow | 30480 | https://www.inaturalist.org/places/geometry/30480.kml |

**Supplementary Table 3:** Renaming and grouping of iNaturalist iconic taxon names into common names.

| **iNaturalist Iconic Taxon Name** | **Common Name** |
|---|---|
| Reptilia | Reptiles and Amphibians |
| Amphibia | |
| Chromista | Protozoa and Chromista |
| Protozoa | |
| Animalia | Other Animalia |

| | |
|---|---|
| Actinopterygii | Fishes |
| Aves | Birds |
| Plantae | Plants |
| Fungi | Fungus |
| Insecta | Insects |
| Arachnida | Spiders |
| Mollusca | Molluscs |
| Mammalia | Mammals |

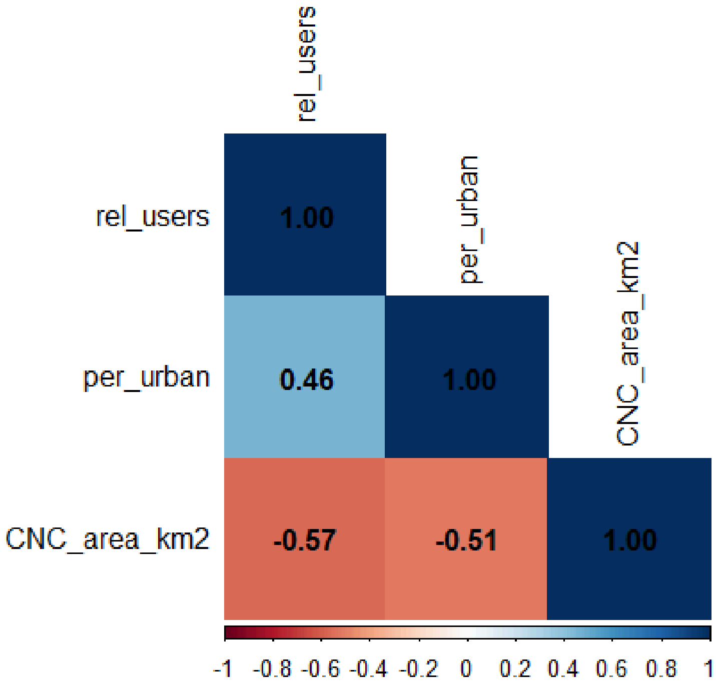


**Supplementary Figure 1:** Pair-wise correlations for predictors used in nb-GLM (per_urban = percent urbanness, rel_users = relative user increase, CNC_area_km2 = project size).

**Supplementary Table 4:** Land cover groupings used in analysis.

| **Original Land Covers** | **Land Cover Grouping for Analysis** |
|---|---|
| Fresh Water, Reeds, Salt Marsh, Foreshore, Riparian, Tidal Water | Blue Space |
| Garden, Allotment, Crops, Grass, Rough Grass, Trees | Green Space |
| Impervious Surface, Buildings, Roads | Grey Space |

**Supplementary Table 5:** Annual participation of City Nature Challenge projects from 2018–2025 (1 = participated, 0 = no participation).

| CNCProject | 2018 | 2019 | 2020 | 2021 | 2022 | 2023 | 2024 | 2025 |
|---|---|---|---|---|---|---|---|---|
| Birmingham and Black Country | 0 | 0 | 1 | 1 | 1 | 1 | 1 | 1 |
| Brighton and Eastern Downs | 0 | 0 | 0 | 1 | 1 | 1 | 1 | 1 |
| Bristol and Bath | 1 | 1 | 1 | 1 | 1 | 1 | 1 | 1 |
| Greater Edinburgh | 0 | 0 | 0 | 1 | 1 | 1 | 1 | 1 |
| Greater Glasgow | 0 | 0 | 0 | 1 | 1 | 1 | 1 | 1 |
| Greater Manchester | 0 | 1 | 1 | 1 | 1 | 1 | 1 | 1 |
| Liverpool City Region | 0 | 1 | 1 | 1 | 1 | 1 | 1 | 1 |
| London | 1 | 1 | 1 | 1 | 1 | 1 | 1 | 1 |
| North East England | 0 | 1 | 1 | 1 | 1 | 1 | 1 | 1 |
| Nottingham City | 0 | 0 | 1 | 1 | 1 | 1 | 1 | 1 |
| Cardiff and Newport | 0 | 0 | 0 | 1 | 1 | 1 | 0 | 1 |
| Coventry | 0 | 0 | 0 | 1 | 1 | 1 | 1 | 0 |
| Gloucestershire | 0 | 0 | 0 | 1 | 1 | 1 | 1 | 0 |
| Lancashire | 0 | 0 | 1 | 1 | 1 | 0 | 1 | 1 |
| Chester Region | 0 | 0 | 0 | 0 | 0 | 1 | 1 | 1 |
| Dundee | 0 | 0 | 0 | 0 | 0 | 1 | 1 | 1 |
| Hull | 0 | 0 | 0 | 0 | 0 | 1 | 1 | 1 |
| Staffordshire & Stoke-On-Trent | 0 | 0 | 0 | 0 | 0 | 1 | 1 | 1 |
| Swansea | 0 | 0 | 0 | 0 | 0 | 1 | 1 | 1 |
| Berkshire | 0 | 0 | 0 | 0 | 0 | 0 | 1 | 1 |

| | | | | | | | | |
|---|---|---|---|---|---|---|---|---|
| Greater Belfast | 0 | 0 | 0 | 0 | 0 | 0 | 1 | 1 |
| Oxfordshire | 0 | 0 | 0 | 0 | 0 | 0 | 1 | 1 |
| Sufflok | 0 | 0 | 0 | 0 | 0 | 1 | 1 | 0 |
| Warwickshire and Solihull | 0 | 0 | 0 | 0 | 0 | 1 | 1 | 0 |
| York | 0 | 0 | 0 | 0 | 0 | 0 | 1 | 1 |
| Congleton | 0 | 0 | 0 | 0 | 0 | 0 | 0 | 1 |
| Derby | 0 | 0 | 0 | 0 | 0 | 1 | 0 | 0 |
| Glamorgan | 0 | 0 | 0 | 0 | 0 | 0 | 0 | 1 |
| Hereforshire | 0 | 0 | 0 | 0 | 0 | 0 | 0 | 1 |
| High Peak | 0 | 0 | 0 | 0 | 0 | 0 | 0 | 1 |
| Oxforshire and Berkshire | 0 | 0 | 0 | 0 | 0 | 1 | 0 | 0 |
| Warwickshire, Solihull and Conventry | 0 | 0 | 0 | 0 | 0 | 0 | 0 | 1 |

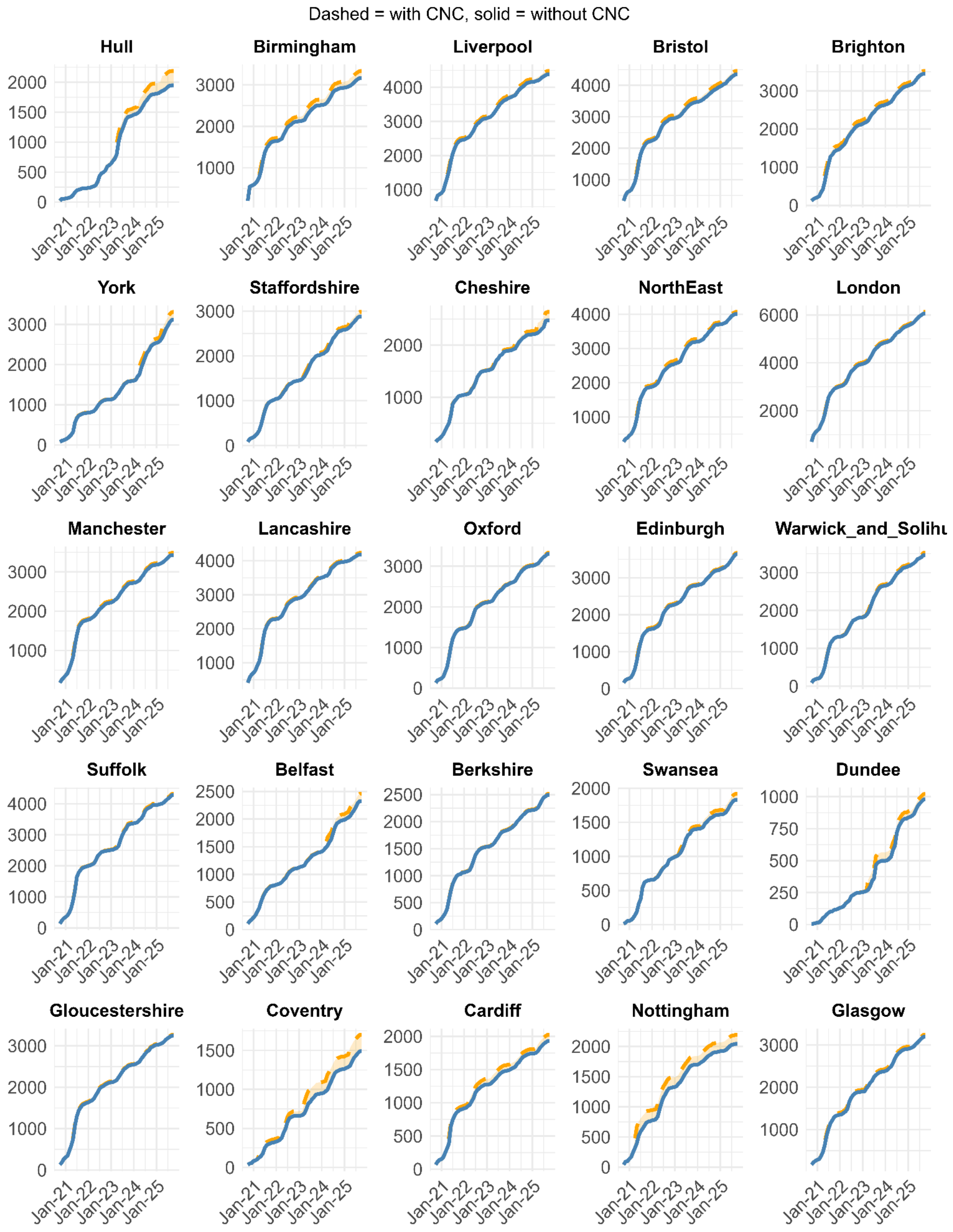


**Supplementary Figure 2:** Species accumulation curves for all projects. The orange dashed line indicates contributions from CNC observations.

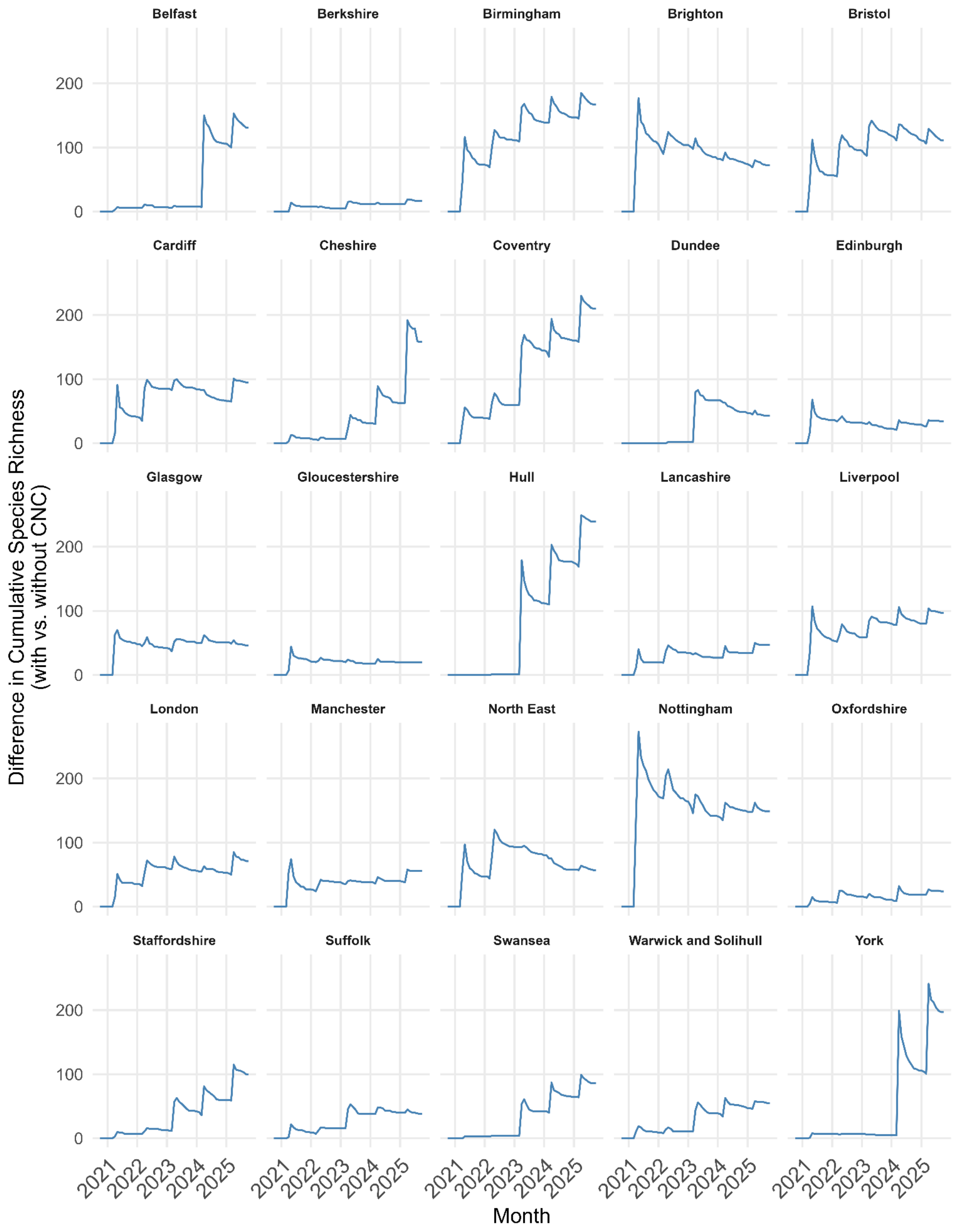


**Supplementary Figure 3:** Difference curves showing the change in cumulative species richness with and without CNC events across projects.

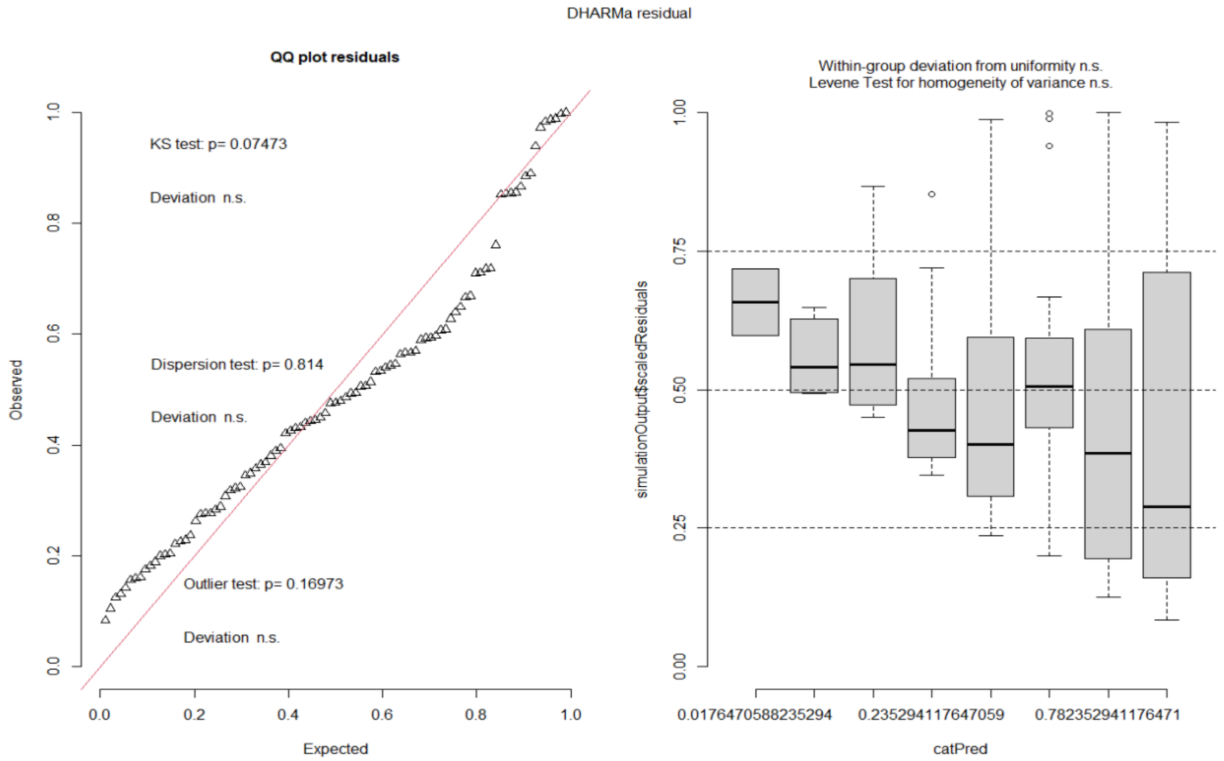


**Supplementary Figure 4:** DHARMa simulated residual diagnostics for the LMM, indicating no major deviations from model assumptions.

**Supplementary Table 6:** LMM model summary (Formula: percent_gain ~ cnc_participation_number + (1 | city))

**Fixed effects**

| Term | Estimate (β) | SE | df | t | p |
|---|---|---|---|---|---|
| Intercept | 8.081 | 1.130 | 54.54 | 7.15 | <0.001 *** |
| CNC participation number | −0.978 | 0.240 | 87.68 | −4.08 | <0.001 *** |

**Random effects**

| Group | Effect | Variance | SD |
|---|---|---|---|
| city | Intercept | 16.093 | 4.012 |
| Residual | — | 9.074 | 3.012 |

**Supplementary Table 7:** Summary of CNC effects on annual species richness across successive years of participation for all 25 CNC projects.

| CNC Year | Mean (%) | SD (%) | Min (%) | Max (%) | Sample Size |
|---|---|---|---|---|---|
| 1 | 5.55 | 4.39 | 0.71 | 15.94 | 17 |
| 2 | 5.76 | 7.24 | 0.74 | 33.83 | 20 |
| 3 | 5.73 | 4.49 | 0.83 | 15.68 | 17 |
| 4 | 4.19 | 3.70 | 0.71 | 13.07 | 14 |
| 5 | 2.91 | 2.30 | 0.85 | 8.30 | 11 |
| 6 | 3.47 | 2.33 | 1.32 | 7.67 | 7 |
| 7 | 1.94 | 0.87 | 1.11 | 3.34 | 5 |
| 8 | 2.13 | 1.07 | 1.37 | 2.89 | 2 |

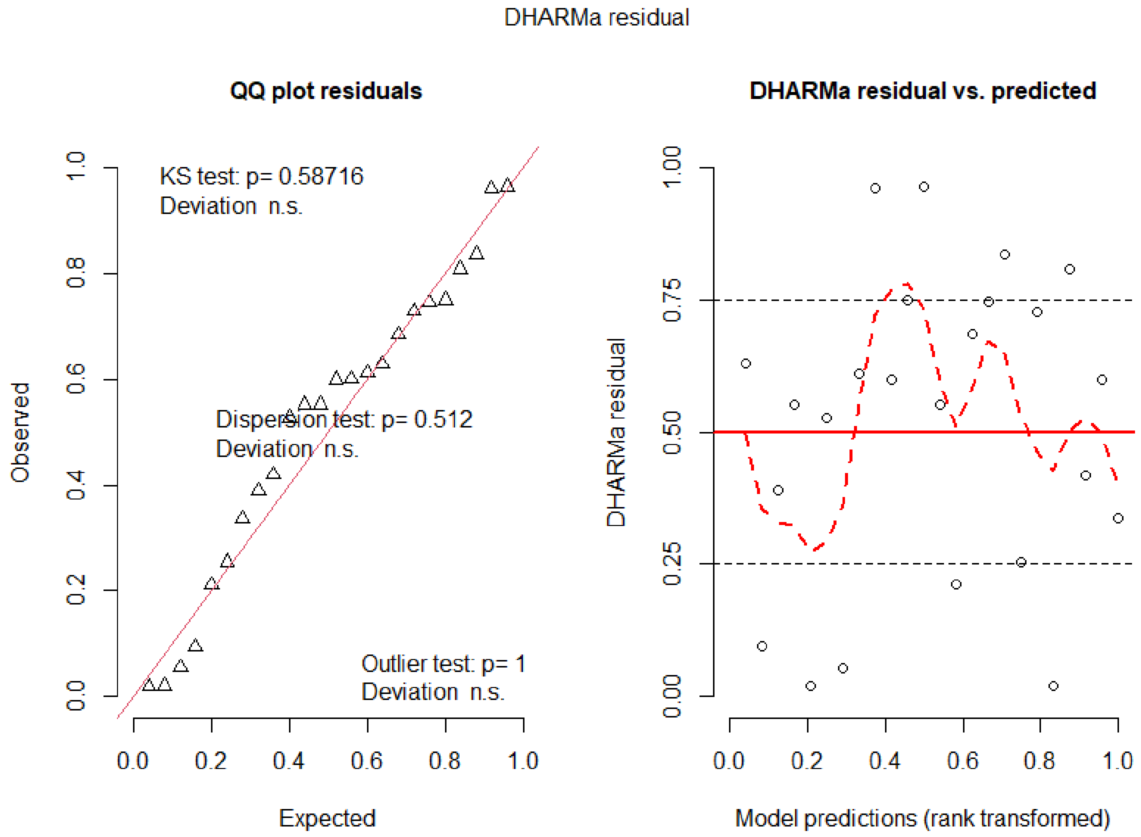


**Supplementary Figure 6:** DHARMa simulated residual diagnostics for the nb-GLM, indicating no major deviations from model assumptions.

**Supplementary Table 8:** nb-GLM model summary (Formula: total_n ~ rel_users + per_urban + CNC_area_km2).

**Coefficients**

| Term | Estimate (β) | SE | z | p |
|---|---|---|---|---|
| Intercept | 1.965 | 0.498 | 3.95 | <0.001 *** |
| rel_users | 1.150 | 0.232 | 4.95 | <0.001 *** |
| per_urban | 0.006 | 0.004 | 1.54 | 0.125 |
| CNC_area_km2 | 0.000017 | 0.000043 | 0.40 | 0.689 |

**Supplementary Table 9:** Composition of unique taxa by taxonomic group for each city.

| | taxon_group | n | percentage | city |
|---|---|---|---|---|
| 1 | Insects | 110 | 46 | Hull |
| 2 | Plants | 75 | 31.4 | Hull |
| 3 | Fungus | 21 | 8.8 | Hull |
| 4 | Other Animalia | 9 | 3.8 | Hull |
| 5 | Molluscs | 6 | 2.5 | Hull |
| 6 | Spiders | 6 | 2.5 | Hull |
| 7 | Protozoa and Chromista | 4 | 1.7 | Hull |
| 8 | Fishes | 3 | 1.3 | Hull |
| 9 | Mammals | 3 | 1.3 | Hull |
| 10 | Birds | 1 | 0.4 | Hull |
| 11 | Insects | 72 | 43.1 | Birmingham |
| 12 | Plants | 48 | 28.7 | Birmingham |
| 13 | Fungus | 18 | 10.8 | Birmingham |
| 14 | Spiders | 11 | 6.6 | Birmingham |
| 15 | Other Animalia | 8 | 4.8 | Birmingham |
| 16 | Molluscs | 4 | 2.4 | Birmingham |
| 17 | Mammals | 3 | 1.8 | Birmingham |
| 18 | Birds | 2 | 1.2 | Birmingham |
| 19 | Reptiles and Amphibians | 1 | 0.6 | Birmingham |
| 20 | Plants | 33 | 34 | Liverpool |

| | | | | |
|---|---|---|---|---|
| 21 | Insects | 32 | 33 | Liverpool |
| 22 | Fungus | 14 | 14.4 | Liverpool |
| 23 | Spiders | 6 | 6.2 | Liverpool |
| 24 | Other Animalia | 5 | 5.2 | Liverpool |
| 25 | Molluscs | 3 | 3.1 | Liverpool |
| 26 | Birds | 2 | 2.1 | Liverpool |
| 27 | Mammals | 1 | 1 | Liverpool |
| 28 | Protozoa and Chromista | 1 | 1 | Liverpool |
| 29 | Insects | 37 | 33.3 | Bristol |
| 30 | Plants | 35 | 31.5 | Bristol |
| 31 | Fungus | 18 | 16.2 | Bristol |
| 32 | Molluscs | 9 | 8.1 | Bristol |
| 33 | Spiders | 5 | 4.5 | Bristol |
| 34 | Other Animalia | 4 | 3.6 | Bristol |
| 35 | Birds | 1 | 0.9 | Bristol |
| 36 | Fishes | 1 | 0.9 | Bristol |
| 37 | Protozoa and Chromista | 1 | 0.9 | Bristol |
| 38 | Plants | 25 | 34.7 | Brighton |
| 39 | Insects | 21 | 29.2 | Brighton |
| 40 | Spiders | 8 | 11.1 | Brighton |
| 41 | Fungus | 7 | 9.7 | Brighton |
| 42 | Molluscs | 5 | 6.9 | Brighton |
| 43 | Birds | 3 | 4.2 | Brighton |

| | | | | |
|---|---|---|---|---|
| 44 | Other Animalia | 3 | 4.2 | Brighton |
| 45 | Insects | 44 | 44 | Staffordshire |
| 46 | Plants | 31 | 31 | Staffordshire |
| 47 | Fungus | 11 | 11 | Staffordshire |
| 48 | Molluscs | 5 | 5 | Staffordshire |
| 49 | Birds | 4 | 4 | Staffordshire |
| 50 | Spiders | 3 | 3 | Staffordshire |
| 51 | Other Animalia | 2 | 2 | Staffordshire |
| 52 | Plants | 85 | 53.8 | Cheshire |
| 53 | Insects | 37 | 23.4 | Cheshire |
| 54 | Fungus | 18 | 11.4 | Cheshire |
| 55 | Spiders | 6 | 3.8 | Cheshire |
| 56 | Other Animalia | 4 | 2.5 | Cheshire |
| 57 | Mammals | 3 | 1.9 | Cheshire |
| 58 | Molluscs | 3 | 1.9 | Cheshire |
| 59 | Birds | 1 | 0.6 | Cheshire |
| 60 | Insects | 93 | 47.2 | York |
| 61 | Plants | 48 | 24.4 | York |
| 62 | Fungus | 24 | 12.2 | York |
| 63 | Other Animalia | 14 | 7.1 | York |
| 64 | Spiders | 11 | 5.6 | York |
| 65 | Fishes | 2 | 1 | York |
| 66 | Molluscs | 2 | 1 | York |
| 67 | Birds | 1 | 0.5 | York |

| | | | | |
|---|---|---|---|---|
| 68 | Mammals | 1 | 0.5 | York |
| 69 | Protozoa and Chromista | 1 | 0.5 | York |
| 70 | Insects | 50 | 70.4 | London |
| 71 | Plants | 8 | 11.3 | London |
| 72 | Other Animalia | 5 | 7 | London |
| 73 | Fungus | 3 | 4.2 | London |
| 74 | Spiders | 2 | 2.8 | London |
| 75 | Birds | 1 | 1.4 | London |
| 76 | Molluscs | 1 | 1.4 | London |
| 77 | Protozoa and Chromista | 1 | 1.4 | London |
| 78 | Other Animalia | 15 | 26.3 | North East |
| 79 | Insects | 13 | 22.8 | North East |
| 80 | Plants | 11 | 19.3 | North East |
| 81 | Molluscs | 8 | 14 | North East |
| 82 | Fungus | 5 | 8.8 | North East |
| 83 | Fishes | 2 | 3.5 | North East |
| 84 | Spiders | 2 | 3.5 | North East |
| 85 | Birds | 1 | 1.8 | North East |
| 86 | Plants | 21 | 37.5 | Manchester |
| 87 | Insects | 20 | 35.7 | Manchester |
| 88 | Fungus | 6 | 10.7 | Manchester |
| 89 | Spiders | 3 | 5.4 | Manchester |
| 90 | Birds | 2 | 3.6 | Manchester |

| | | | | |
|---|---|---|---|---|
| 91 | Other Animalia | 2 | 3.6 | Manchester |
| 92 | Fishes | 1 | 1.8 | Manchester |
| 93 | Reptiles and Amphibians | 1 | 1.8 | Manchester |
| 94 | Plants | 25 | 53.2 | Lancashire |
| 95 | Insects | 11 | 23.4 | Lancashire |
| 96 | Fungus | 4 | 8.5 | Lancashire |
| 97 | Molluscs | 3 | 6.4 | Lancashire |
| 98 | Spiders | 2 | 4.3 | Lancashire |
| 99 | Other Animalia | 1 | 2.1 | Lancashire |
| 100 | Protozoa and Chromista | 1 | 2.1 | Lancashire |
| 101 | Insects | 8 | 33.3 | Oxfordshire |
| 102 | Plants | 8 | 33.3 | Oxfordshire |
| 103 | Other Animalia | 4 | 16.7 | Oxfordshire |
| 104 | Fungus | 2 | 8.3 | Oxfordshire |
| 105 | Mammals | 1 | 4.2 | Oxfordshire |
| 106 | Protozoa and Chromista | 1 | 4.2 | Oxfordshire |
| 107 | Plants | 16 | 47.1 | Edinburgh |
| 108 | Insects | 9 | 26.5 | Edinburgh |
| 109 | Other Animalia | 3 | 8.8 | Edinburgh |
| 110 | Birds | 2 | 5.9 | Edinburgh |
| 111 | Fungus | 2 | 5.9 | Edinburgh |
| 112 | Molluscs | 1 | 2.9 | Edinburgh |
| 113 | Spiders | 1 | 2.9 | Edinburgh |

| | | | | |
|---|---|---|---|---|
| 114 | Insects | 26 | 47.3 | Warwick and Solihull |
| 115 | Fungus | 14 | 25.5 | Warwick and Solihull |
| 116 | Plants | 11 | 20 | Warwick and Solihull |
| 117 | Spiders | 2 | 3.6 | Warwick and Solihull |
| 118 | Mammals | 1 | 1.8 | Warwick and Solihull |
| 119 | Plants | 13 | 34.2 | Suffolk |
| 120 | Insects | 12 | 31.6 | Suffolk |
| 121 | Spiders | 5 | 13.2 | Suffolk |
| 122 | Fungus | 4 | 10.5 | Suffolk |
| 123 | Molluscs | 2 | 5.3 | Suffolk |
| 124 | Other Animalia | 2 | 5.3 | Suffolk |
| 125 | Plants | 53 | 40.5 | Belfast |
| 126 | Fungus | 19 | 14.5 | Belfast |
| 127 | Insects | 18 | 13.7 | Belfast |
| 128 | Other Animalia | 14 | 10.7 | Belfast |
| 129 | Molluscs | 13 | 9.9 | Belfast |
| 130 | Protozoa and Chromista | 5 | 3.8 | Belfast |
| 131 | Mammals | 4 | 3.1 | Belfast |
| 132 | Spiders | 2 | 1.5 | Belfast |
| 133 | Birds | 1 | 0.8 | Belfast |
| 134 | Insects | 9 | 52.9 | Berkshire |

| | | | | |
|---|---|---|---|---|
| 135 | Plants | 6 | 35.3 | Berkshire |
| 136 | Birds | 2 | 11.8 | Berkshire |
| 137 | Plants | 40 | 46.5 | Swansea |
| 138 | Insects | 27 | 31.4 | Swansea |
| 139 | Fungus | 5 | 5.8 | Swansea |
| 140 | Birds | 4 | 4.7 | Swansea |
| 141 | Spiders | 4 | 4.7 | Swansea |
| 142 | Mammals | 2 | 2.3 | Swansea |
| 143 | Other Animalia | 2 | 2.3 | Swansea |
| 144 | Molluscs | 1 | 1.2 | Swansea |
| 145 | Protozoa and Chromista | 1 | 1.2 | Swansea |
| 146 | Plants | 30 | 69.8 | Dundee |
| 147 | Insects | 4 | 9.3 | Dundee |
| 148 | Fungus | 3 | 7 | Dundee |
| 149 | Molluscs | 2 | 4.7 | Dundee |
| 150 | Protozoa and Chromista | 2 | 4.7 | Dundee |
| 151 | Other Animalia | 1 | 2.3 | Dundee |
| 152 | Spiders | 1 | 2.3 | Dundee |
| 153 | Insects | 9 | 45 | Gloucestershire |
| 154 | Plants | 8 | 40 | Gloucestershire |
| 155 | Fungus | 2 | 10 | Gloucestershire |
| 156 | Molluscs | 1 | 5 | Gloucestershire |
| 157 | Plants | 98 | 46.7 | Coventry |

| | | | | |
|---|---|---|---|---|
| 158 | Insects | 72 | 34.3 | Coventry |
| 159 | Fungus | 19 | 9 | Coventry |
| 160 | Molluscs | 7 | 3.3 | Coventry |
| 161 | Other Animalia | 4 | 1.9 | Coventry |
| 162 | Birds | 3 | 1.4 | Coventry |
| 163 | Spiders | 3 | 1.4 | Coventry |
| 164 | Fishes | 1 | 0.5 | Coventry |
| 165 | Mammals | 1 | 0.5 | Coventry |
| 166 | Protozoa and Chromista | 1 | 0.5 | Coventry |
| 167 | Plants | 36 | 37.9 | Cardiff |
| 168 | Insects | 26 | 27.4 | Cardiff |
| 169 | Molluscs | 9 | 9.5 | Cardiff |
| 170 | Fungus | 6 | 6.3 | Cardiff |
| 171 | Mammals | 5 | 5.3 | Cardiff |
| 172 | Other Animalia | 5 | 5.3 | Cardiff |
| 173 | Spiders | 4 | 4.2 | Cardiff |
| 174 | Birds | 2 | 2.1 | Cardiff |
| 175 | Fishes | 1 | 1.1 | Cardiff |
| 176 | Reptiles and Amphibians | 1 | 1.1 | Cardiff |
| 177 | Plants | 72 | 48.3 | Nottingham |
| 178 | Insects | 50 | 33.6 | Nottingham |
| 179 | Other Animalia | 8 | 5.4 | Nottingham |
| 180 | Spiders | 7 | 4.7 | Nottingham |

| | | | | |
|---|---|---|---|---|
| 181 | Fungus | 5 | 3.4 | Nottingham |
| 182 | Molluscs | 5 | 3.4 | Nottingham |
| 183 | Birds | 2 | 1.3 | Nottingham |
| 184 | Insects | 20 | 43.5 | Glasgow |
| 185 | Plants | 12 | 26.1 | Glasgow |
| 186 | Fungus | 4 | 8.7 | Glasgow |
| 187 | Molluscs | 4 | 8.7 | Glasgow |
| 188 | Other Animalia | 3 | 6.5 | Glasgow |
| 189 | Birds | 1 | 2.2 | Glasgow |
| 190 | Fishes | 1 | 2.2 | Glasgow |
| 191 | Spiders | 1 | 2.2 | Glasgow |

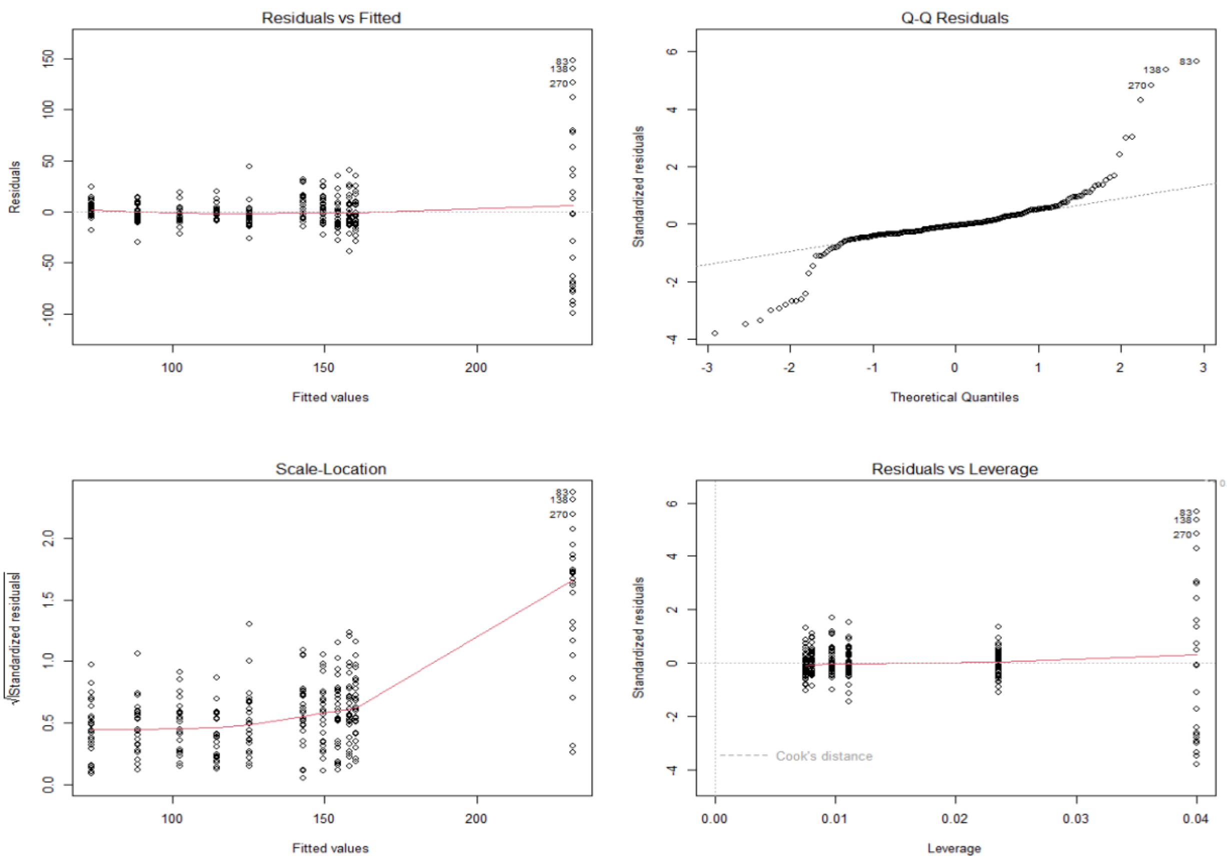


**Supplementary Figure 7:** Standard diagnostic plots for the linear model. Diagnostics indicate acceptable model fit, with some increased variance at higher fitted values consistent with the large response values associated with CNC event effects.

**Supplementary Table 10:** Model summary for linear model (Formula = rel_users ~ is_cnc_week + poly(weeks_from_cnc, 2).

**Fixed effects**

| Term | Estimate (β) | SE | t | p |
|---|---|---|---|---|
| Intercept | 127.782 | 1.695 | 75.39 | <0.001 *** |
| CNC week | 96.533 | 5.981 | 16.14 | <0.001 *** |
| Weeks from CNC (poly 1) | 455.761 | 26.621 | 17.12 | <0.001 *** |
| Weeks from CNC (poly 2) | −105.220 | 28.512 | −3.69 | <0.001 *** |

**Supplementary Table 11:** Pairwise bootstrap comparisons of land-cover metrics among grid activity types (n = 141 grids per group; 1,000 bootstrap replicates).

| comparison | median_diff | lower_CI | upper_CI | p_value | metric | sig |
|---|---|---|---|---|---|---|
| CNC-only - Active | 0.12361831 | 0.02865507 | 0.2278256 | 0.007 | Blue | Significant |
| CNC-only - Active | -0.1103688 | -0.19672899 | -0.0329586 | 0.993 | Shannon | Significant |
| CNC-only - Inactive | 0.21483049 | 0.13518881 | 0.2964717 | 0 | Blue | Significant |
| Active - Inactive | 0.08432037 | 0.040459 | 0.1365437 | 0 | Blue | Significant |
| Active - Inactive | 0.0557727 | 0.01188802 | 0.1032982 | 0.01 | Shannon | Significant |
| CNC-only - Active | -0.01335305 | -0.07416003 | 0.0497063 | 0.658 | Green | Not significant |
| CNC-only - Active | -0.03252798 | -0.08805578 | 0.0284557 | 0.863 | Grey | Not significant |
| CNC-only - Inactive | -0.03189434 | -0.09418907 | 0.026119 | 0.857 | Green | Not significant |
| CNC-only - Inactive | -0.04755169 | -0.10561391 | 0.0057077 | 0.959 | Grey | Not significant |
| CNC-only - Inactive | -0.05247947 | -0.14786346 | 0.0335649 | 0.889 | Shannon | Not significant |
| Active - Inactive | -0.01867942 | -0.05405718 | 0.0140569 | 0.847 | Green | Not significant |

| | | | | | | |
|---|---|---|---|---|---|---|
| Active - Inactive | -0.01796299 | -0.05064649 | 0.0160406 | 0.837 | Grey | Not significant |
